# Estimating DNA methylation potential energy landscapes from nanopore sequencing data

**DOI:** 10.1101/2021.02.22.431480

**Authors:** Jordi Abante, Sandeep Kambhampati, Andrew P. Feinberg, John Goutsias

**Affiliations:** Whitaker Biomedical Engineering Institute, Johns Hopkins University, Baltimore, MD 21218, USA; Department of Electrical and Computer Engineering, Johns Hopkins University, Baltimore, MD 21218, USA; Department of Biomedical Engineering, Johns Hopkins University, Baltimore, MD 21218 USA; Center for Epigenetics, Johns Hopkins University School of Medicine, Baltimore, MD 21205, USA; Department of Medicine, Johns Hopkins University School of Medicine, Baltimore, MD 21205, USA

## Abstract

High-throughput third-generation sequencing devices, such as the Oxford Nanopore Technologies (ONT) MinION sequencer, can generate long reads that span thousands of bases. This new technology opens the possibility of considering a wide range of epigenetic modifications and provides the capability of interrogating previously inaccessible regions of the genome, such as highly repetitive regions, as well as of performing comprehensive allele-specific methylation analysis, among other applications. It is well-known, however, that detection of DNA methylation from nanopore data results in a substantially reduced per-read accuracy when comparing to WGBS, due to noise introduced by the sequencer and its underlying chemistry. It is therefore imperative that methods are developed for the reliable modeling and analysis of the DNA methylation landscape using nanopore data. Here we introduce such method that takes into account the presence of noise introduced by the ONT sequencer and, by using simulations, we provide evidence of its potential. The proposed approach establishes a solid foundation for the development of a comprehensive framework for the statistical analysis of DNA methylation, and possibly of other epigenetic marks, using third-generation sequencing.

## Introduction

DNA methylation through 5-methylcytosine (5mC) is an important biochemical process that influences biological function in cells by establishing stable and inheritable epigenetic marks throughout the genome^1^. By using a sodium bisulfite treatment and second generation sequencing, whole-genome bisulfite sequencing (WGBS) generates methylation profiles with comprehensive genomic coverage, high quantitative accuracy, and excellent reproducibility^2^. However, WGBS produces short methylation reads with low contextual information, which limits the scope and effectiveness of downstream analysis^3^.

High-throughput third-generation sequencing devices, such as the Oxford Nanopore Technologies (ONT) MinION sequencer, can generate long reads that span thousands of bases. This is facilitated by detecting methylation marks from electrolytic signals produced by these devices using available methylation calling software, such as Nanopolish^4^, DeepSignal^5^, DeepMod^6^, and Megalodon^7^, and provides a decisive edge over short-read bisulfite sequencing in a number of appealing biological applications^8–10^. It is well-known, however, that detection of DNA methylation from nanopore sequencing data produces substantially lower per-read accuracy than WGBS due to noise introduced by the sequencer and its underlying chemistry.

Nanopolish, for example, uses a versatile method of signal detection theory^11^ based on a log-likelihood ratio approach to detect 5mC methylation in nanopore reads. This is achieved by calling a CpG site to be methylated when the associated log-likelihood ratio exceeds a given threshold *T >* 0, and to be unmethylated when this ratio is below −*T*. Although this approach is known to maximize the true positive rate for a fixed value of the false positive rate, it has several drawbacks. For example, even though accuracy (i.e., the probability that a CpG site is correctly predicted to be methylated or unmethylated) can be improved by increasing *T*, this comes at the expense of making fewer calls. Indeed, analysis by Simpson et al.^4^ (see Supplementary Figure 2, pore model R9) suggests that to achieve an accuracy of 95% using Nanopolish, about 32% of the nanopore-based calls must be discarded, although this could be even larger in reality.

In addition, and by focusing on a small set of 100 CpG sites in the *E. coli* reference genome using a fully methylated and fully unmethylated control dataset from PCR-amplified DNA and M.SssI-treated DNA, Yuen et al.^12^ have recently provided evidence of deficient detection performance for a number of nanopore-based methylation callers, including the ones mentioned above. These callers exhibited a trade-off between true positive and false positive rates and produced poor empirical estimates of the true methylation mean values.

Notably, comprehensive downstream statistical analysis of the methylation landscape requires knowledge of first- and higher-order methylation statistics along the genome, such as means, correlations, methylation entropies, and probability distributions^13–16^. Unfortunately, these quantities cannot be computed reliably from the output of available methylation callers, such as Nanopolish, due to their inferior error rate performance and limited data coverage. Therefore, new statistical approaches must be developed to address the previous issues by taking into account the distinctive nature of nanopore sequencing^3, 9^.

We here introduce a novel approach that characterizes the probability distribution of methylation within a genomic region of interest using a parametric correlated potential energy landscape (CPEL) model that is consistent with methylation means and pairwise correlations at each CpG site. To fit this model to methylation data obtained from nanopore sequencing, we propose an estimation approach based on the expectation-maximization (EM) algorithm^17^. This method determines values for the parameters of the CPEL model by maximizing the likelihood that the observed nanopore sequencing data have been generated by the estimated model.

To demonstrate the potential of our approach for modeling nanopore methylation data, we evaluated its performance by employing partially methylated synthetic data, which were generated by a simulation-based approach that we designed using human WGBS and nanopore sequencing data. By employing these data, we evaluated Nanopolish with a method that is more realistic than the ones used by Simpson et al.^4^ and Yuen et al.^12^, which were based on fully methylated and fully unmethylated data. Our results show that Nanopolish can achieve reasonable per-read detection performance at low levels of nanopore signal noise, which substantially deteriorates at higher noise levels common in nanopore sequencing. Importantly, our simulations revealed poor error rate performance in calling the true values of the sufficient statistics^18^ of our model, providing compelling evidence that reliable estimation of methylation statistics from nanopore data requires the development of a method that takes into account the presence of noise introduced by the ONT sequencer.

Using simulations, we also evaluated our approach to modeling DNA methylation landscapes from nanopore sequencing data. Our preliminary results show the potential of the proposed EM-based maximum-likelihood method for reliably estimating the parameters of a CPEL model for DNA methylation, even at a low coverage, by employing information produced by processing the raw nanopore signals using Nanopolish. This provides a solid foundation for the development of a comprehensive framework for the statistical analysis of DNA methylation, and potentially other binary epigenetic markers, using third-generation sequencing.

## Modeling DNA methylation stochasticity

For reasons to become clear in the following section, we follow Nanopolish and cluster the CpG sites along the reference genome into groups, so that the last C nucleotide of a group and the first C nucleotide of the next group are separated by at least 10 bases. We then form the base sequence between the first C nucleotide and the last G nucleotide in each group and extend each sequence by 5 bases upstream and 4 bases downstream (see Fig. 1a). This process defines DNA segments 𝒢_*l*_, *l* = 1, 2, …, along the genome, which we refer to as *CG-groups*, that include at least one CpG site. We found that 84.1% (18,770,638) of the CG-groups in the human reference genome (autosomal chromosomes, built GRCh38.p12) contain 11 bases, and therefore include 1 CpG site, whereas the remaining 15.9% (3,549,239) of the CG-groups contain from 12 to 1002 bases and include 2 to 212 CpG sites.

**Figure 1.**
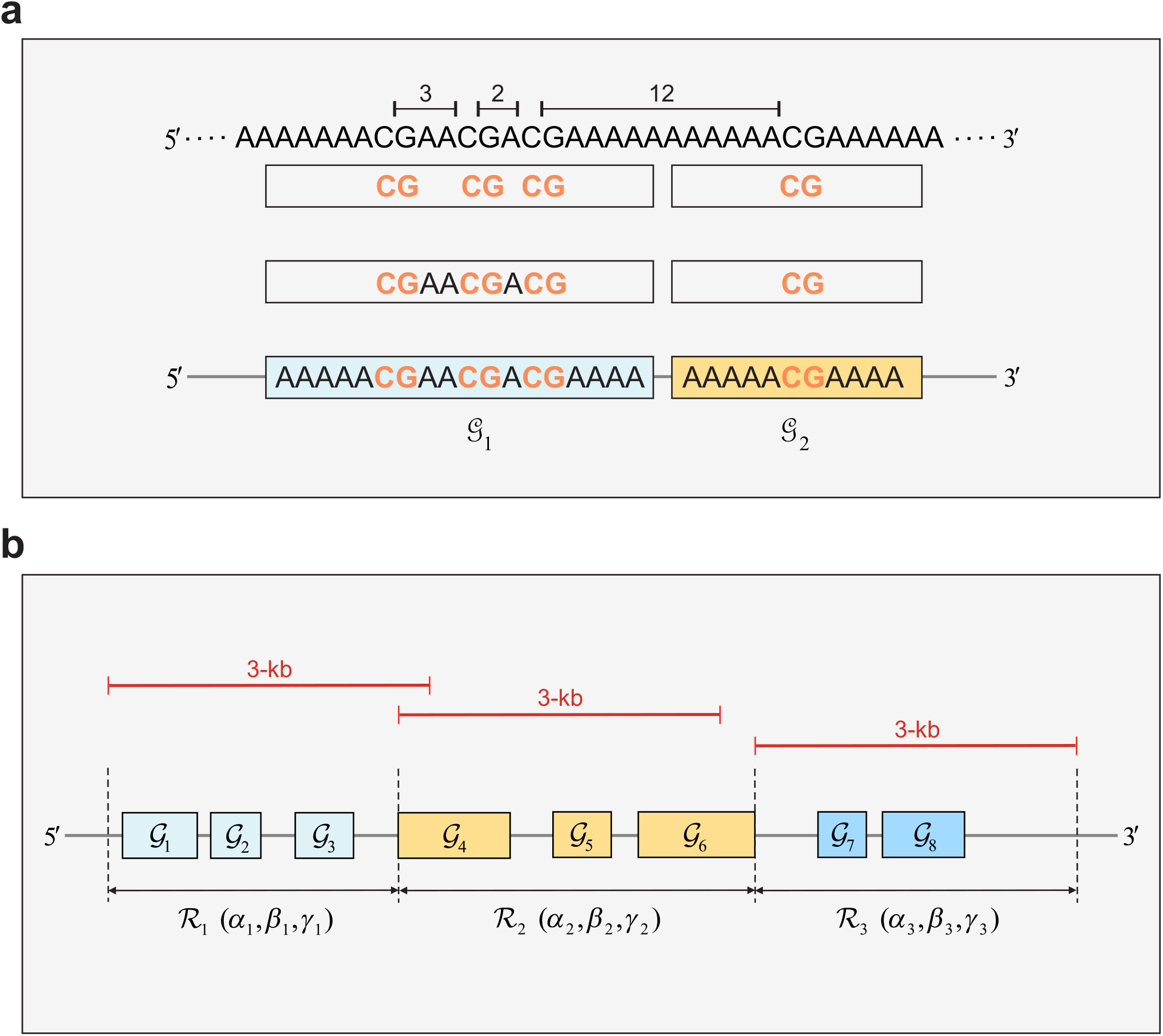
CG-groups and modeling regions. **a** Example of clustering CpG sites along the genome into groups of CG dinucleotides that are separated by at least 10 bases. The depicted CG-groups 𝒢_1_ and 𝒢_2_ are formed from two CG clusters by inserting the bases between the first C nucleotide and the last G nucleotide in each cluster and by extending each sequence by 5 bases upstream and 4 bases downstream. **b** Example of partitioning a given chromosome into modeling regions ℜ. Region ℜ_1_ is less than 3000 bases long, whereas ℜ_2_ is more than 3000 bases long. This guarantees that each modeling region fully contains CG-groups. Note that the CPEL model given by Eqs. (7)-(9) is associated with different parameters *α, β*, and *γ* within each modeling region, whose values must be estimated using nanopore data corresponding to the specific region.

We also partition each chromosome into non-overlapping genomic regions ℛ_*k*_, *k* = 1, 2, …, which we refer to as *modeling regions*, using the following recursive scheme (see Fig. 1b for an example). For *k* = 0, 1, …, we place a 3-kb window along the genome starting at the first nucleotide after the most downstream nucleotide in ℛ_*k*_ (when *k* = 0, we take the starting nucleotide to be the first nucleotide in the chromosome). This window defines the modeling region *R*_*k*+1_, provided that its most downstream nucleotide does not intersect a CG-group. Note, however, that if this is not true and if the window contains more than 50% of an intersecting CG-group, then we increase its size to form the smallest region ℛ_*k*+1_ that fully contains the intersecting CG-group. Otherwise, we decrease the window to form the largest region ℛ_*k*+1_ whose downstream nucleotide does not intersect a CG-group. We found 66 modeling regions ℛ in the human genome (of size between 2994 and 3000 bases) containing 1 CG-group, which include 1 or 2 CpG sites each, and 919,255 modeling regions (of size between 2905 and 3071 bases) containing at least 2 CG-groups, which include from 2 to 487 CpG sites each.

Within a modeling region ℛ that contains *N* CpG sites *n* = 1, 2, …, *N*, we characterize DNA methylation using the *N* × 1 random methylation state vector ***X*** = [*X*_1_ *X*_2_ … *X*_*N*_]^*T*^, where *X*_*n*_ = 0, if the *n*-th CpG site is unmethylated, and *X*_*n*_ = 1, if the CpG site is methylated. As a consequence of the well-known maximum-entropy principle^19^, the probability distribution of methylation (PDM) that is consistent with methylation means and pairwise correlations at each CpG site is given by

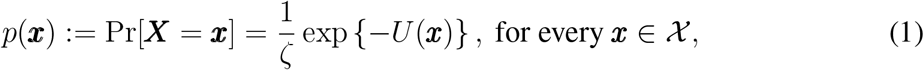

where χ is the set of all 2^*N*^ possible methylation patterns associated with ℛ,

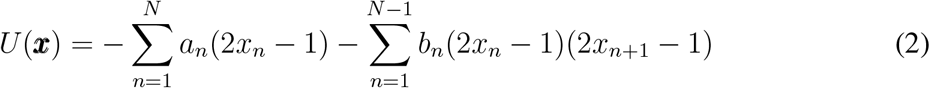

is the potential energy function of ***X***, *a*_*n*_ and *b*_*n*_ are two parameters associated with the *n*-th CpG site, and

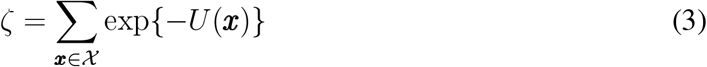

is a normalizing constant known as the partition function. Note that parameter *a*_*n*_ affects the propensity of the *n*-th CpG site to be methylated without the influence of nearby CpG sites, whereas parameter *b*_*n*_ accounts for the possibility that the methylation states of two contiguous CpG sites *n* and *n* + 1 would be correlated.

The previous PDM generalizes the well-known one-dimensional Ising model of statistical physics^20^ by including ‘external field’ parameters *a*_*n*_, *n* = 1, 2, …, *N*, and ‘interaction’ parameters *b*_*n*_, *n* = 1, 2, …, *N* − 1, which are not necessarily constant. Note however that this PDM does not account for evidence suggesting that the likelihood of a given CpG site to be methylated depends strongly on the fraction of CpG sites within a local neighborhood, as well as on the methylation status of nearby CpG sites whose influence diminishes as their nucleotide distance from the given CpG site increases^21, 22^. To address this issue, we follow the approach by Jenkinson et al.^13, 14^ and set

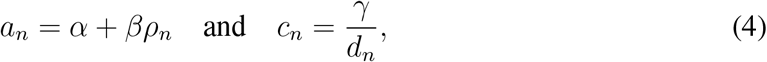

where *α, β*, and *γ* are parameters characteristic to the modeling region ℛ, *ρ*_*n*_ is the CpG density, defined as the fraction of dinucleotides that are CpG sites within a symmetric neighborhood of 1000 nucleotides centered at the *n*-th CpG site, given by

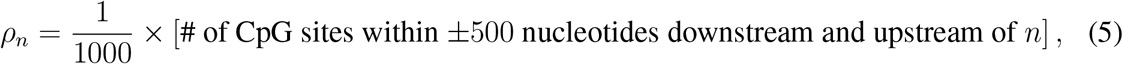

and *d*_*n*_ is the distance of the *n*-th CpG site from its downstream CpG site *n* + 1, given by

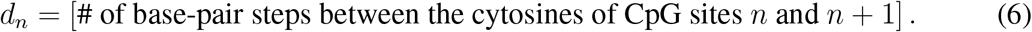

Notably, parameter *α* accounts for intrinsic factors that affect the propensity of CpG sites to be methylated, whereas parameters *β* and *γ* respectively modulate the influence of CpG density and distance on methylation, which we assume here to be applied uniformly on all CpG sites within each modeling region ℛ.

As a consequence of Eqs. (1)-(6), we characterize the methylation vector ***X*** within a modeling region ℛ that contains *N* CpG sites *n* = 1, 2, …, *N* using the PDM

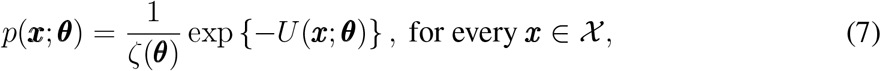

with potential energy function

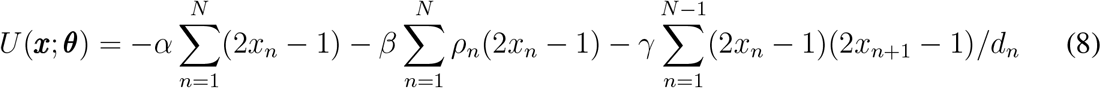

and partition function

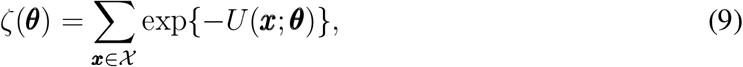

where ***θ*** = [*α β γ*]^*T*^ is the vector of the underlying parameters associated with region ℛ. We refer to this PDM as the correlated potential energy landscape (CPEL) model. Notably, this model summarizes the common understanding that methylation of a CpG site depends on two distinct factors: the local CpG architecture, quantified by CpG densities and distances, as well as the biochemical environment provided by the methylation machinery, quantified by parameters *α, β*, and *γ*. Moreover, Eqs. (7)-(9) lead to an Ising model that has been successful in predicting the probability of DNA methylation within regions of the genome from CpG density and distance alone^13^. Note however that additional sequence-specific factors may influence DNA methylation^23, 24^, which could be included in future versions of the model if necessary.

## Maximum-likelihood parameter estimation

To use the previous CPEL model for methylation analysis of nanopore sequencing data, we need to estimate its parameters ***θ*** using data that contain sufficient information about the true methylation state so that the inferred PDM best predicts the statistical behavior of these data within a modeling region ℛ. Performing this task requires the availability of a set ***y*** of nanopore observations related to the methylation state ***x*** of ℛ. Given such observations, we can then find a vector 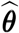 that maximizes the likelihood that the observed nanopore data ***y*** have been generated by the CPEL model with parameters 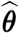 by solving the following optimization problem:

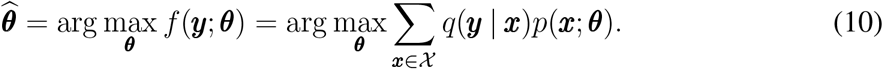

In this equation, *q*(***y*** | ***x***) is the conditional probability distribution of observable nanopore data ***Y***, given that the methylation state in ℛ is ***x***, and *f* (***y***; ***θ***) is the marginal probability distribution of ***Y*** when the parameter values of the generating CPEL model are given by ***θ***. Notably, both *p* and *q* depend on the genetic context. For notational convenience, however, we do not explicitly denote this dependence here.

To proceed with maximum-likelihood parameter estimation, we need to identify appropriate observable nanopore data ***Y*** that are informative of the methylation state ***X*** of ℛand compute the conditional probability distribution *q*(***y*** | ***x***). We do so by employing the hidden Markov model (HMM) of Nanopolish. Although a number of alternative approaches have been recently proposed in the literature for addressing this problem using artificial neural networks, including DeepMod, DeepSignal, and Megalodon, only Nanopolish can be used to produce the conditional probability distribution *q*(***y*** | ***x***), which is essential for inferring a presumed stochastic DNA methylation model from available nanopore data.

To explain how this is done, first note that a nucleotide in a given DNA fragment remains inside a pore of the nanopore sequencer for some time during which the instrument continuously samples the electrical current flow generated across the pore. Analysis of the resulting raw nanopore samples by a step detection algorithm can be employed to identify abrupt changes in the current signal during sequencing that define ‘events’ of relatively stationary signal behavior indicating discrete motion of the DNA sequence through the pore. Each event can then be characterized by its duration, as well as by the average and dispersion of the current values measured during the event, three statistical summaries that can be used to perform base-calling^4, 25^.

Nanopolish uses an appropriately designed and trained hidden Markov model (HMM) to identify the sequence of nanopore events associated with a given nanopore read, determine the methylation status of each CpG site in the read, and map this information to the human reference genome. For a given genomic region ℛ that fully contains a set {𝒢_*l*_, *l* = 1, 2, …, *L*} of CG-groups, Nanopolish evaluates the conditional probability distribution *q*_*l*_(***y***_*l*_ | ***x***_*l*_) of the vector ***Y*** _*l*_ of all average nanopore current values mapped to a CG-group 𝒢_*l*_(***x***_*l*_) whose CpG sites are methylated or unmethylated in accordance to the corresponding methylation state ***x***_*l*_. By setting ***Y*** = [***Y*** _1_ ***Y*** _2_ … ***Y*** _*L*_]^*T*^ and by making the reasonable assumption that, given the methylation state ***x*** in ℛ, the averages of the nanopore current values within the individual CG-groups 𝒢_*l*_, *l* = 1, 2, …, *L*, are conditionally independent, we obtain

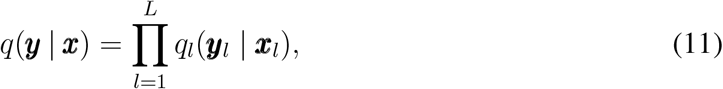

which allows us to compute *q*(***y*** |***x***) using Nanopolish. Due to lack of data, however, Nanopolish cannot in general compute the conditional probability distributions *q*_*l*_ associated with all CG-groups within a genomic region ℛ, in which case our formulas must be appropriately modified to account for this realistic scenario.

Unfortunately, the current version of Nanopolish employs an HMM that has been trained using only fully methylated or fully unmethylated data^4^. Consequently, it can be used in its current form to compute the conditional probability distribution *q*_*l*_(***y***_*l*_ | ***x***_*l*_) only when the values of ***x***_*l*_ are all ones (fully methylated state) or all zeros (fully unmethylated sate). To take into account this issue, note that

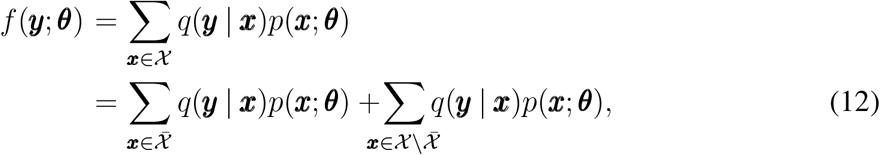

where 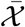 is the set of all methylation states in χ for which the CpG sites within the CG-groups in ℛ are all methylated or unmethylated, and 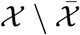 are the remaining states. Then, by considering evidence that DNA methylation over CpG sites that are closely clustered to each other are most often strongly correlated^21, 22^, we can assume that

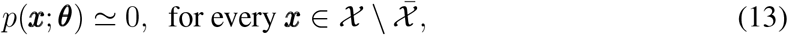

in which case we can approximately set

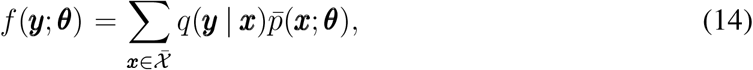

where

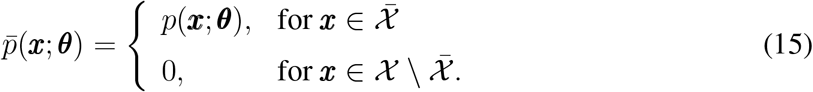

This implies that

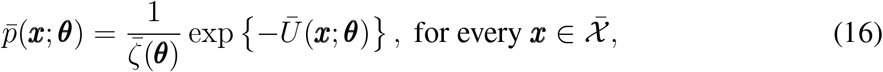

where

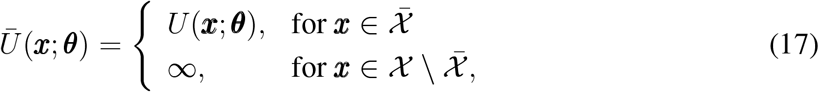

and

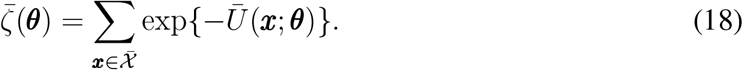

Notably, 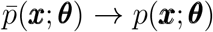 as 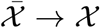, which shows that, if Nanopolish is trained in the future using partially methylated data, then the previous approximation can be eliminated.

With each CG-group G_*l*_, we now associate the random variable

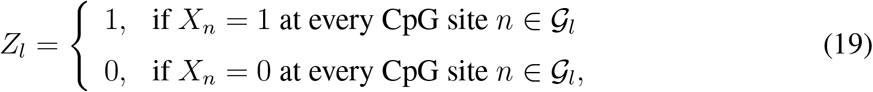

which we refer to as the *methylation index* of the CG-group, since its value indicates whether the CG-group is fully methylated or fully unmethylated. From Eqs. (8), (16), (17), and (19) we can show that, if ***Z*** = [*Z*_1_ *Z*_2_ … *Z*_*L*_]^*T*^ is the random vector of methylation indices associated with the CG-groups within a modeling region ℛ, then its probability distribution *π*(***z***; ***θ***) := Pr[***Z*** = ***z***; ***θ***] is given by

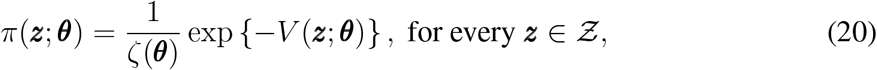

with potential energy function

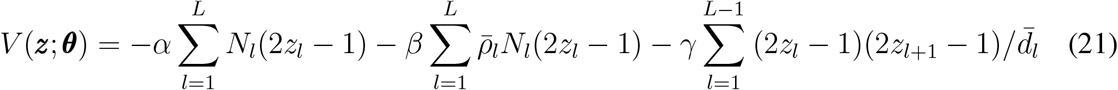

and partition function

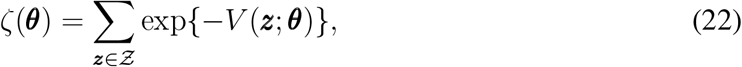

where Ƶ is the set of all possible 2^*L*^ methylation index values in ℛ. In these equations, which we refer to as the *reduced* CPEL model,

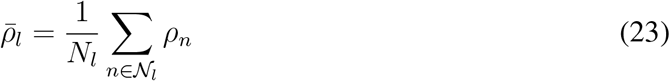

is the average CpG density within the *l*-th CG-group 𝒢_*l*_ containing *N*_*l*_ CpG sites in N_*l*_, and 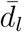 is the distance between the last CpG site in the CG-group 𝒢_*l*_ and the first CpG site in the CG-group 𝒢_*l*+1_. Note that, although the reduced CPEL model always depends on parameters *α* and *β*, it will also depend on the interaction parameter *γ*, provided that ℛ contains at least two CG-groups. In this case, we can estimate the original CPEL model from available Nanopolish data by fitting the reduced CPEL model to that data. For this reason, we do not model regions ℛ that contain only one CG-group, which are nevertheless very few (we found only 66 such regions in the human genome) and are insignificant (each of these regions contain only 1 or 2 CpG sites). In addition, and for reliable parameter estimation, we only model regions that contain at least 10 CpG sites, with average coverage of at least 5× per CG-group, and for which methylation information is available for at least 2*/*3 of their CG-groups.

Note now that, every methylation pattern 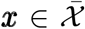 can be generated by a *unique* vector of methylation indices ***z*** ∈ Ƶ. Consequently, if ***x*** = ***s***(***z***) is the methylation pattern in 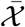 associated with ***z***, then we have from Eq. (14) that

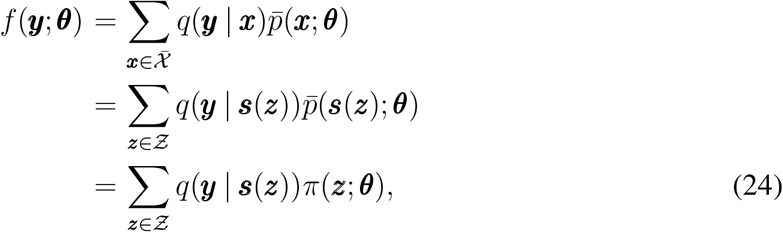

by virtue of the fact that 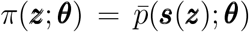. This result, together with Eqs. (10) and (11), leads to the following maximum-likelihood parameter estimation problem:

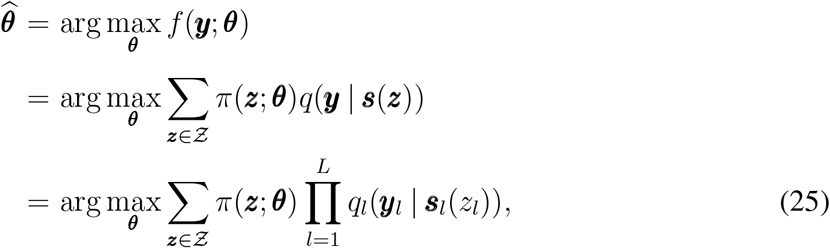

where ***s***_*l*_(*z*_*l*_) is either the fully methylated pattern within 𝒢_*l*_ (when *z*_*l*_ = 1) or the fully unmethylated pattern (when *z*_*l*_ = 0). Solving this problem facilitates estimation of the values of the parameters of the original CPEL model via maximum likelihood using Nanopolish data. An example that summarizes the main steps of this method is depicted in Fig. 2.

**Figure 2.**
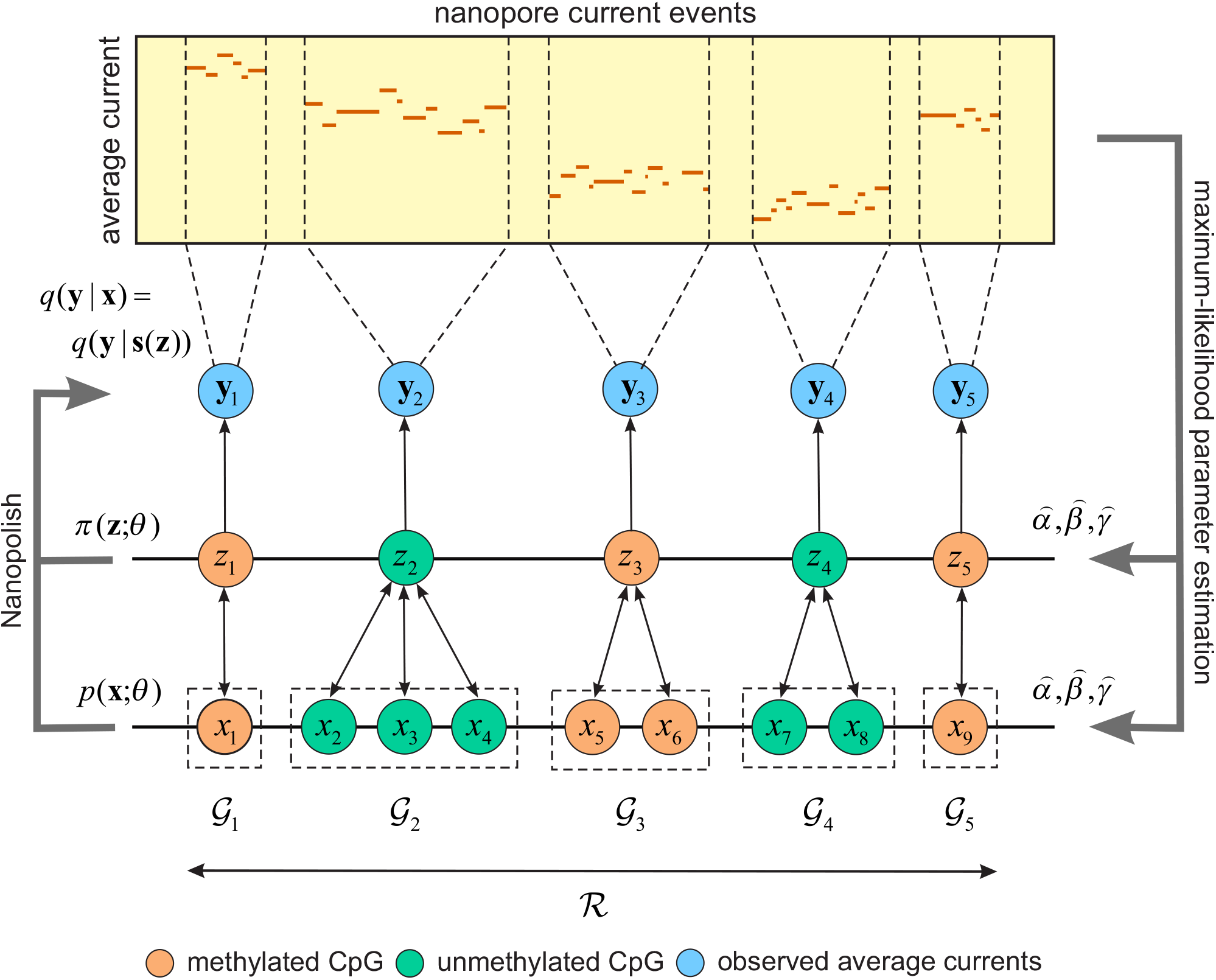
Maximum-likelihood estimation of the CPEL model. Within a modeling region R that fully contains five CG-groups 𝒢_1_, 𝒢_2_, 𝒢_3_, 𝒢_4_, and 𝒢_5_, the parameters *α, β*, and *γ* of the CPEL model *p*(***x***; ***θ***) are estimated by maximizing the likelihood that observed average current events ***y*** = {***y***_1_, ***y***_2_, ***y***_3_, ***y***_4_, ***y***_5_} (blue) have been generated by the reduced CPEL model *π*(***z***; ***θ***), where ***z*** = {*z*_1_, *z*_2_, *z*_3_, *z*_4_, *z*_5_} are the methylation indices associated with the CG-groups. This requires knowledge of the conditional probability distribution *q*(***y*** | ***x***) = *q*(***y*** | ***s***(***z***)) of the observed average current events given the methylation state ***x*** = ***s***(***z***) within ℜ in which all CpG sites within each CG-group are either methylated (orange) or unmethylated (green), which is computed by Nanopolish.

## Performing parameter estimation using the EM algorithm

In the previous parameter estimation approach, we need to consider the availability of multiple independent nanopore reads 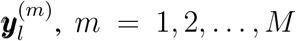, associated with each CG-group 𝒢_*l*_, in which case, we must solve the following maximum-likelihood problem:

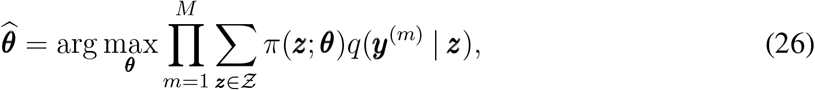

where

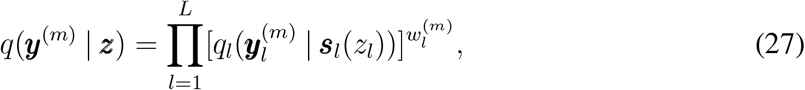

with 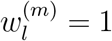, if the *m*-th observation is present in 𝒢_*l*_, and 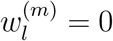, if this observation is missing. Notably, and due to the required summation over all possible methylation indices ***z*** within a modeling region ℛ, evaluating the likelihood function in Eq. (26) is not computationally feasible when ℛ contains many CG-groups and, therefore, directly performing maximum-likelihood parameter estimation using this equation is not appropriate. However, we can address this problem by employing the expectation-maximization (EM) algorithm which results in iteratively applying the following two steps:

### Expectation step

Given nanopore data ***y*** = {***y***^(1)^, ***y***^(2)^, …, ***y***^(*M*)^} and a currently estimated value 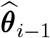 of the parameters ***θ***, compute the conditional expected value of the logarithm of the likelihood function *p*(***y, Z***) with respect to the methylation index vector ***Z***, given by

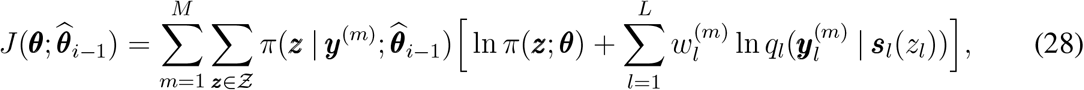

where 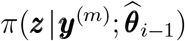 is the posterior probability distribution of the methylation index vector ***Z*** given the nanopore data ***y***^(*m*)^.

### Maximization step

Given 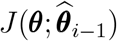, find a new parameter estimate 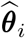 by

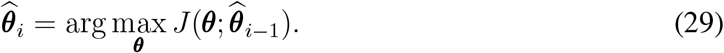

To implement the expectation step, note from Eqs. (20) and (21) that

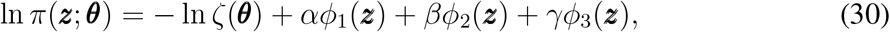

where ***θ*** = [*α β γ*]^*T*^,

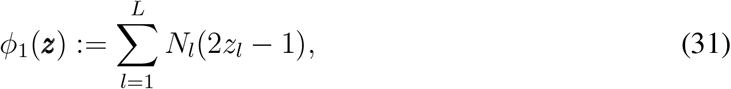

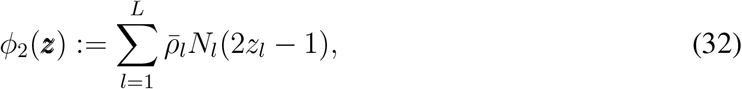

and

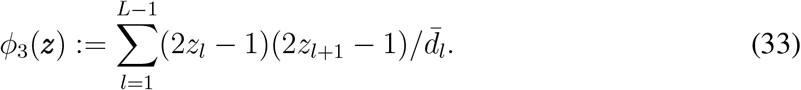

Consequently, Eq. (28) becomes

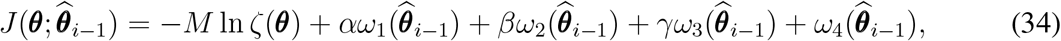

where

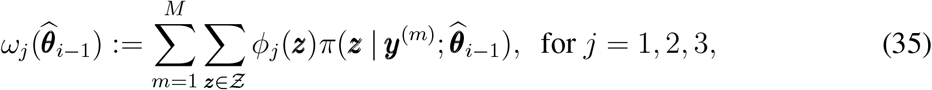

and

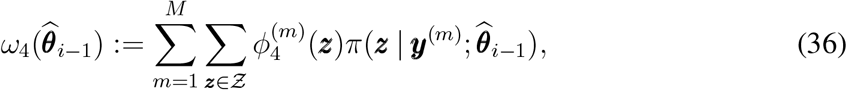

with

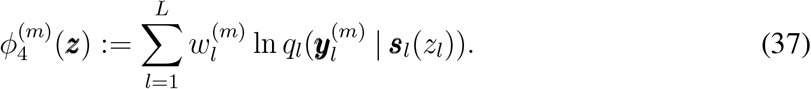

To implement the maximization step, we set the gradient of 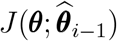 with respect to ***θ*** = [*α β γ*]^*T*^ equal to zero at 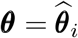. In this case, and from Eqs. (21), (22), and (31)-(34), we obtain the following system of nonlinear equations

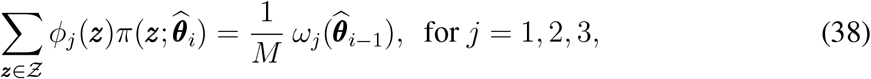

which we must solve to obtain the new parameter value 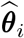.

The previous EM algorithm requires evaluation of the posterior probability distribution *π*(***z*** | ***y***; ***θ***) of the methylation index vector ***Z***, as well as evaluation of expectations and correlations of ***Z*** with respect to its prior and posterior probability distributions. We discuss these computations in the Supplementary Methods. Moreover, we need to solve Eqs. (38), which we carry out by using NLsolve, a Julia implementation of the trust region approach^26^.

## Results

Since the proposed method for modeling methylation stochasticity from nanopore data relies on Nanopolish, we first evaluated the detection performance of this methylation caller by employing a simulation-based benchmarking approach, which we designed using human WGBS and nanopore sequencing data (see Fig. 3). The performance of Nanopolish has been previously investigated^4, 12^ using a small number of CpG sites and a real control dataset from PCR-amplified DNA and M.SssI-treated DNA, with the first comprising fully unmethylated CpG sites and the second fully methylated sites. Our benchmarking method however allows us to evaluate Nanopolish over an entire chromosome using simulated DNA fragments that are not necessarily fully methylated or unmethylated. By using this approach, we obtained results similar to the ones previously obtained using real data, thus demonstrating its legitimacy as a convenient and inexpensive computational tool for evaluating the performance of methylation callers. Moreover, and in addition to providing evidence that the statistical properties of DNA methylation cannot be reliably inferred from the output of Nanopolish, we also show that this can be achieved by using a CPEL model whose parameters can be estimated via EM-based maximum-likelihood from information provided by Nanopolish.

**Figure 3.**
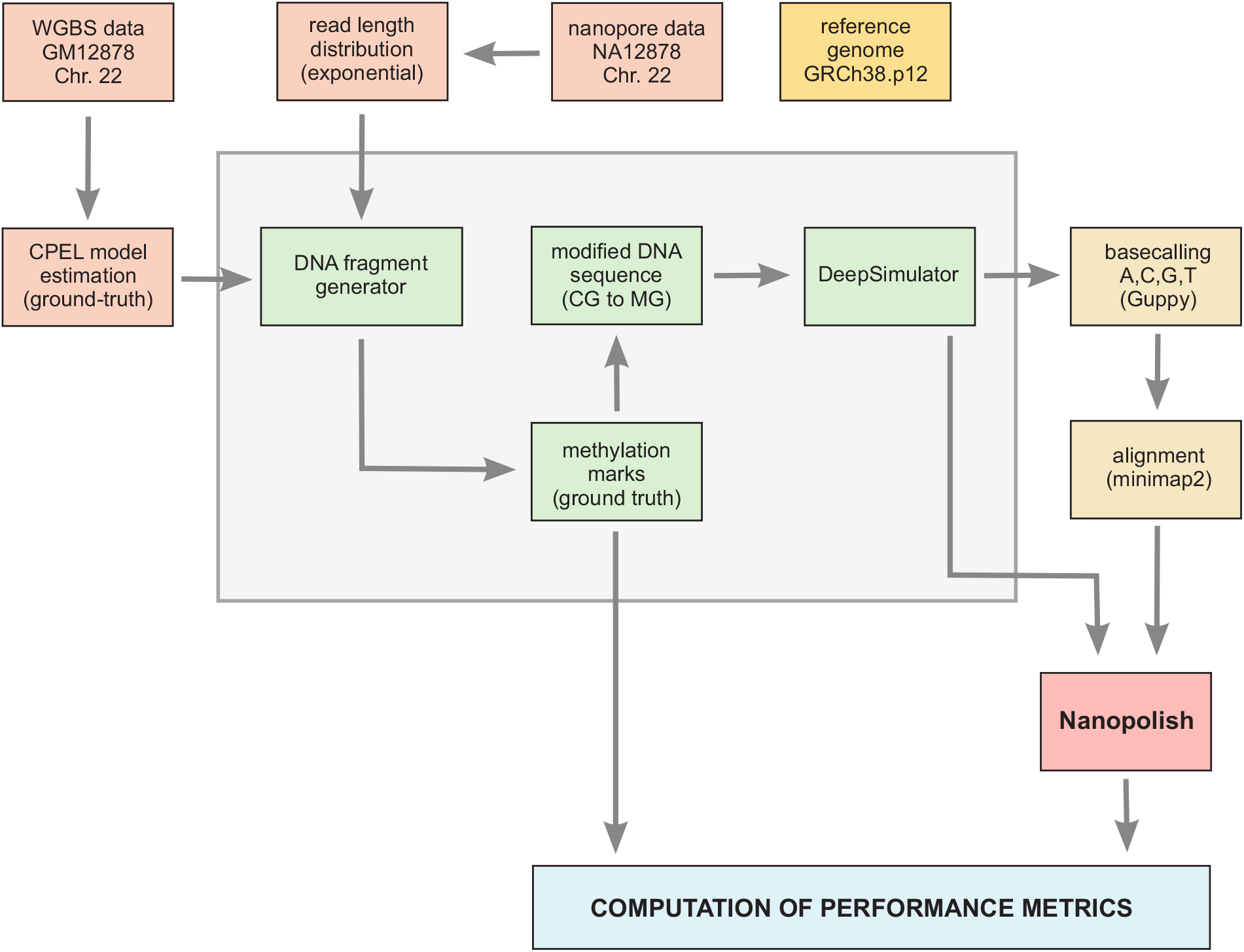
A scheme for benchmarking Nanopolish. A simulation-based benchmarking approach for evaluating the methylation calling performance of Nanopolish. This approach uses human cell-line WGBS and nanopore sequencing data to generate DNA fragments of known methylation states, which are then processed by the DeepSimulator to produce realistic nanopore reads. Evaluation is performed by comparing the output of the caller to the known ground truth methylation states of the input DNA fragments that generate these reads.

### Performance evaluation of Nanopolis

By employing WGBS data associated with the well-characterized human GM12878 Utah/Ceph lymphoblastoid cell line, we used our benchmarking method to construct, within Chr. 22 of the human reference genome that contains 622,083 CpG sites, a ‘ground-truth’ CPEL methylation model given by Eqs. (7)-(9). We did so by estimating, using a previous maximum-likelihood approach^13^, the parameters *α, β*, and *γ* of the model within 3-kb modeling regions ℛ that contain sufficient data to perform reliable estimation (regions that contain at least 10 CpG sites, with an average coverage of at least 5× per CpG site, and for which methylation information is available for at least 2*/*3 of their CpG sites). To determine parameter values within the remaining modeling regions, the method uses all estimated *α, β*, and *γ* values and computes their empirical probability distributions (see Fig. 4). It then assigns parameter values to these regions by drawing samples from the corresponding empirical distributions.

**Figure 4.**
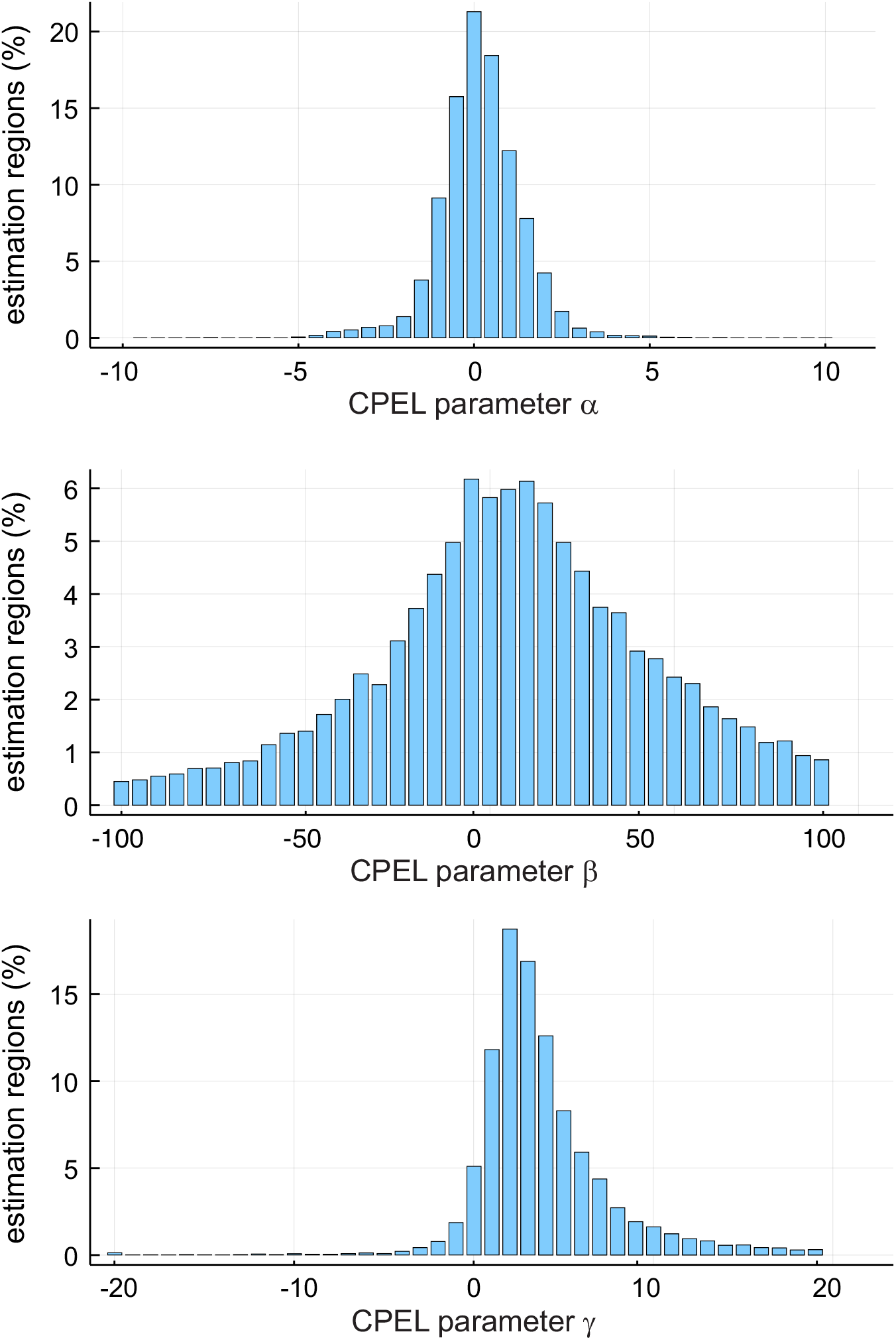
Distributions of CPEL model parameter values. Distributions, within Chr. 22, of the values of the CPEL model parameters *α, β*, and *γ*, which are estimated via maximum-likelihood from human Utah/Ceph lymphoblastoid WGBS data (GM12878).

To generate DNA fragments that satisfy the length distribution and coverage requirements of nanopore reads, our method computes the length distribution of nanopore reads in available nanopore sequencing data associated with the GM12878 cell line^8^. In agreement with Li et al.^27, 28^, we found that the read lengths in these data follow an exponential distribution with rate 1.18 × 10^*−*4^, which we estimated from the real data via maximum-likelihood (see Fig. 5). The method then generates DNA fragments by determining their start location and length along Chr. 22. The start location of a fragment is specified by randomly drawing a number between 1 and *L* − *M* + 1, where *L* = 50,818,468 bp is the length of Chr. 22, whereas its length *M* (in bp) is computed by sampling the previous exponential length distribution. In order to control the methylation coverage, fragment generation is repeated until the average of all nucleotide coverages within Chr. 22 is no less than 25×, with the coverage at each nucleotide being computed as the number of DNA fragments overlapping the nucleotide.

**Figure 5.**
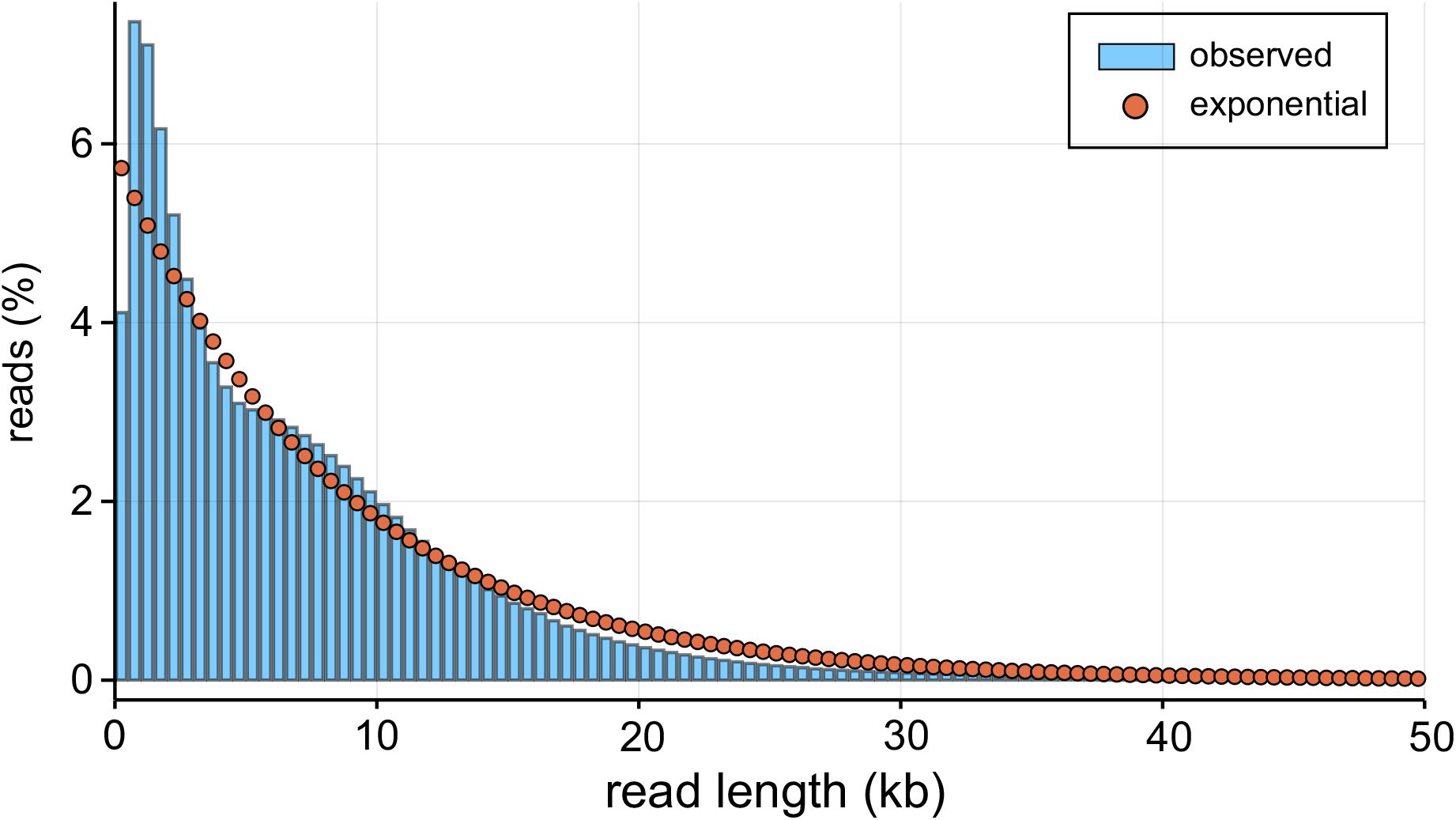
Length distribution of nanopore reads. The distribution of read lengths observed in the GM12878 human nanopore data (blue) was found to be approximately exponential (red dots) with rate 1.18 × 10^*−*4^.

Subsequently, and for each DNA fragment, our benchmarking method identifies all modeling regions that overlap the fragment and, for each modeling region, it generates its methylation state by sampling the ground truth CPEL model associated with that region using a Markov Chain approach (see Supplementary Methods). This information is then used to assign a ground truth binary methylation state to each CpG site in the DNA fragment by marking its CG dinucleotides as being methylated (1) or unmethylated (0) based on their methylation status in the associated modeling regions. Finally, and within each DNA fragment, the C’s of all CG dinucleotide marked by 1 are replaced with M’s, a step that modifies the DNA sequence within each fragment by incorporating the methylation of CG dinucleotides, as determined by the methylation states drawn from the ground truth CPEL model.

Each modified DNA fragment generated by the previous approach was processed by the DeepSimulator^27, 28^, a computational tool that faithfully simulates the entire pipeline of nanopore sequencing and produces nanopore reads consisting of the current values measured by the nanopore. We used the DeepSimulator in its context-independent mode. However, and in order to take into account methylated 6-mers, we replaced its pore model, which is based on ONT’s official statistics, with the one used in Nanopolish. In addition, we generated the raw nanopore reads by adding random noise on the event sequence using the default option of the DeepSimulator and by setting the cutoff frequency of the low-pass filter, which removes high-frequency components from the signal generated from the event sequence, to its default value. Finally, and in order to investigate the effect of signal noise on methylation calling, we added zero-mean Gaussian noise to the raw nanopore reads with standard deviations (sd’s) ranging from 2 to 3.5, which encompasses values normally observed in Nanopolish’s pore model.

The raw nanopore reads produced by the DeepSimulator were used to perform base calling via ONT’s Guppy (CPU mode) whose output was then aligned to the reference genome by minmap2^29^. The aligned data, together with the raw nanopore reads produced by the DeepSimulator, were then fed as inputs to Nanopolish (see Fig. 3). Its output was then used to quantify methylation calling performance by computing several performance metrics, which included accuracy (probability that a CpG site is correctly predicted to be methylated or unmethylated), precision (probability that a CpG site is correctly predicted to be methylated), true positive rate, and true negative rate (see Figs. 6 & 7).

**Figure 6.**
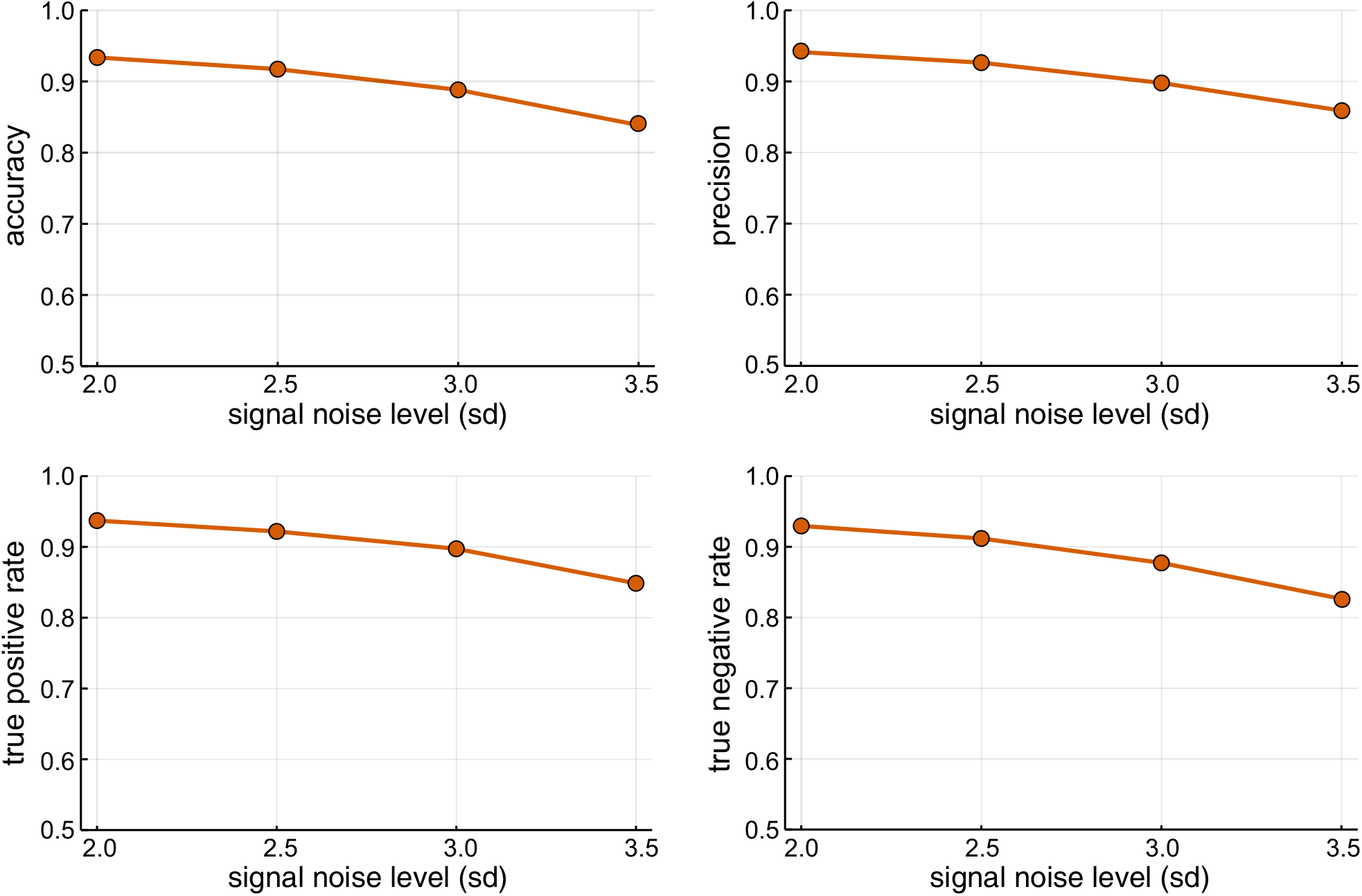
Performance characteristics of Nanopolish. Accuracy, precision, true positive rate, and true negative rate characteristics of Nanopolish at several signal noise levels. The results were obtained by the simulation-based benchmarking method depicted in Fig. 3 and by setting the log-likelihood ratio detection threshold used by Nanopolish equal to zero.

**Figure 7.**
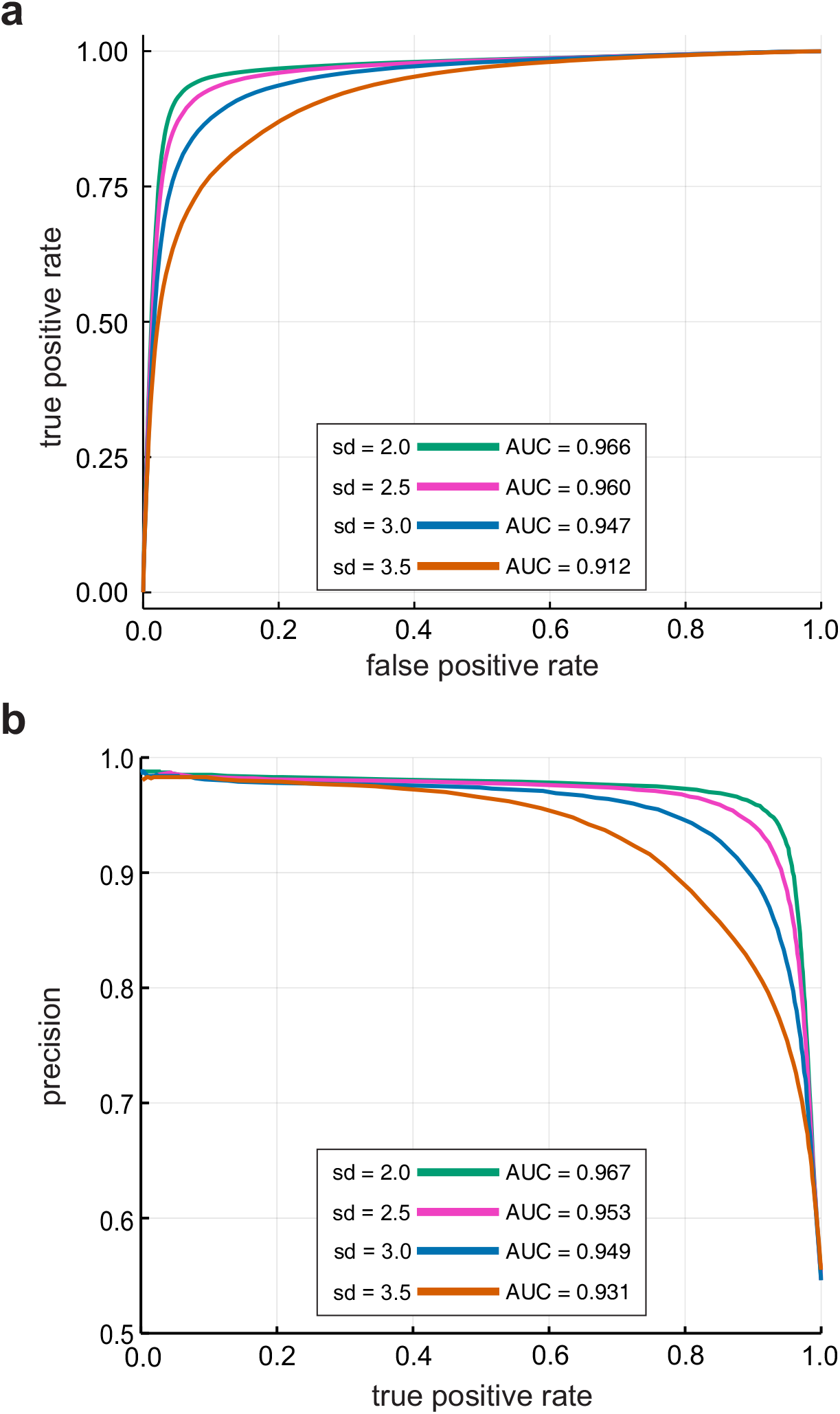
Detection performance of Nanopolish. Receiver operating characteristic curves of Nanopolish, in **a**, and precision-recall curves, in **b**, for several signal noise levels, obtained by the simulation-based benchmarking method depicted in Fig. 3. The area under the curve (AUC) values are also provided in each case.

As expected, Nanopolish exhibited reduced per-read detection performance at higher signal noise levels (see Fig. 6). Moreover, it achieved no more that 94% accuracy, precision, true positive rate, and true negative rate at all noise levels, which dropped to less than 90% for sd ≥ 3. In addition, the receiver operating characteristic (ROC) and precision recall (PR) curves depicted in Fig. 7, calculated by varying the log-likelihood ratio detection threshold used by Nanopolish, showed an expected trade-off between the true positive rate and the false positive rate, as well as a trade-off between precision and true positive rate (also known as recall). Notably, the ROC and PR curves obtained for sd = 3 are similar to those previously reported by Simpson et al.^4^ (Fig. 2, pore R9) and Yuen et al.^12^ (Figs. 3a,b), suggesting that a signal noise level of about 3 is close to reality.

In addition to the previous results, the area under the ROC curve depicted in Fig. 7a decreases with increasing noise, ranging between 0.947 and 0.912 for 3 ≤ sd ≤ 3.5, indicating that Nanopolish exhibits in our simulations only a 91.2% to 94.7% chance of distinguishing between truly methylated and truly unmethylated CpG sites at those noise levels. Notably, using real data, Yuen et al.^12^ reported an area under the ROC curve of 0.921 for Nanopolish. A similar remark is also true for the area under the PR curve depicted in Fig. 7b, which ranges between 0.949 and 0.931 when 3 ≤ sd ≤ 3.5, as compared to the value of 0.924 reported by Yuen et al.^12^ Note that the area under the PR curve can be interpreted as the fraction of true detections made by a randomly selected threshold^30^. We may therefore conclude that, when 3 ≤ sd ≤ 3.5, Nanopolish exhibits only a 93.1% to 94.9% chance that calls at individual CpG sites made by a randomly selected log-likelihood ratio detection threshold will be true methylation calls.

Having established the validity of our simulation-based benchmarking approach and having identified a signal noise level close to 3 as being realistic for base-calling using Nanopolish, we can now ask whether the output of Nanopolish can be directly used for downstream statistical analysis of methylation. To answer this question, we should consider the fact that such an analysis should at least use methylation means at individual CpG sites as well as pairwise correlations at consecutive CpG sites. As previously argued for the case of WGBS data, this necessitates the use of a stochastic model for the methylation state, such as the CPEL model used in this paper, whose parameters must be estimated from available data. We will therefore not be able to answer the previous question in the affirmative if the sufficient statistics associated with the parameters of a given model could not be correctly computed from the methylation calls made by Nanopolish.

The CPEL model given by Eqs. (7)-(9) is a three-parameter exponential family of distributions with (natural) sufficient statistic given by^18^

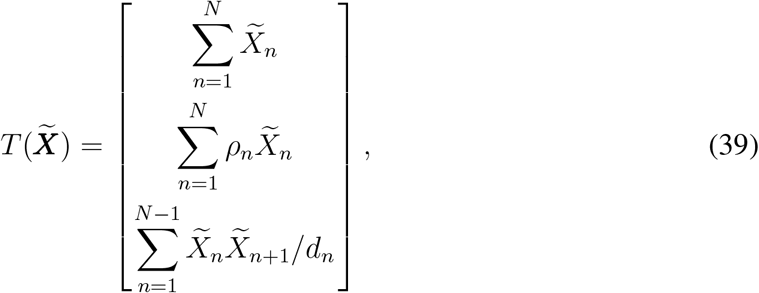

where 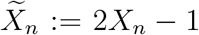. Precise computation of this statistic using Nanopolish depends on its ability to correctly call the true values of 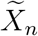 and 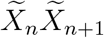, which is directly influenced by the method’s accuracy in detecting methylation at individual CpG sites, as well as in identifying concordant methylation at consecutive CpG sites. Our simulations showed that, if the log-likelihood ratio detection threshold is set to zero, then the error rate (defined as 1 − accuracy) in calling the true value of 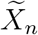 ranges between 11% and 16% when 3 ≤ sd ≤ 3.5 (see Fig. 8a). Notably, this error monotonically decreases to zero when the threshold increases, but this is achieved by substantially reducing the number of methylation calls made by Nanopolish. For example, Fig. 8a shows that, to achieve an error rate of 5% for sd = 3, Nanopolish must produce methylation calls at only 73% of the CpG sites considered, which is in agreement with Simpson et al.^4^ who reported a 6% error rate using a log-likelihood ratio detection threshold of that produced calls at 77% of the targeted CpG sites. This provides evidence that accurate computation of the sufficient statistics 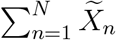 and 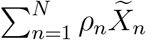, associated with the parameters *α* and *β* of the CPEL model, will require use of a sufficiently high log-likelihood ratio detection threshold, thus resulting in a substantial loss of methylation calls.

**Figure 8.**
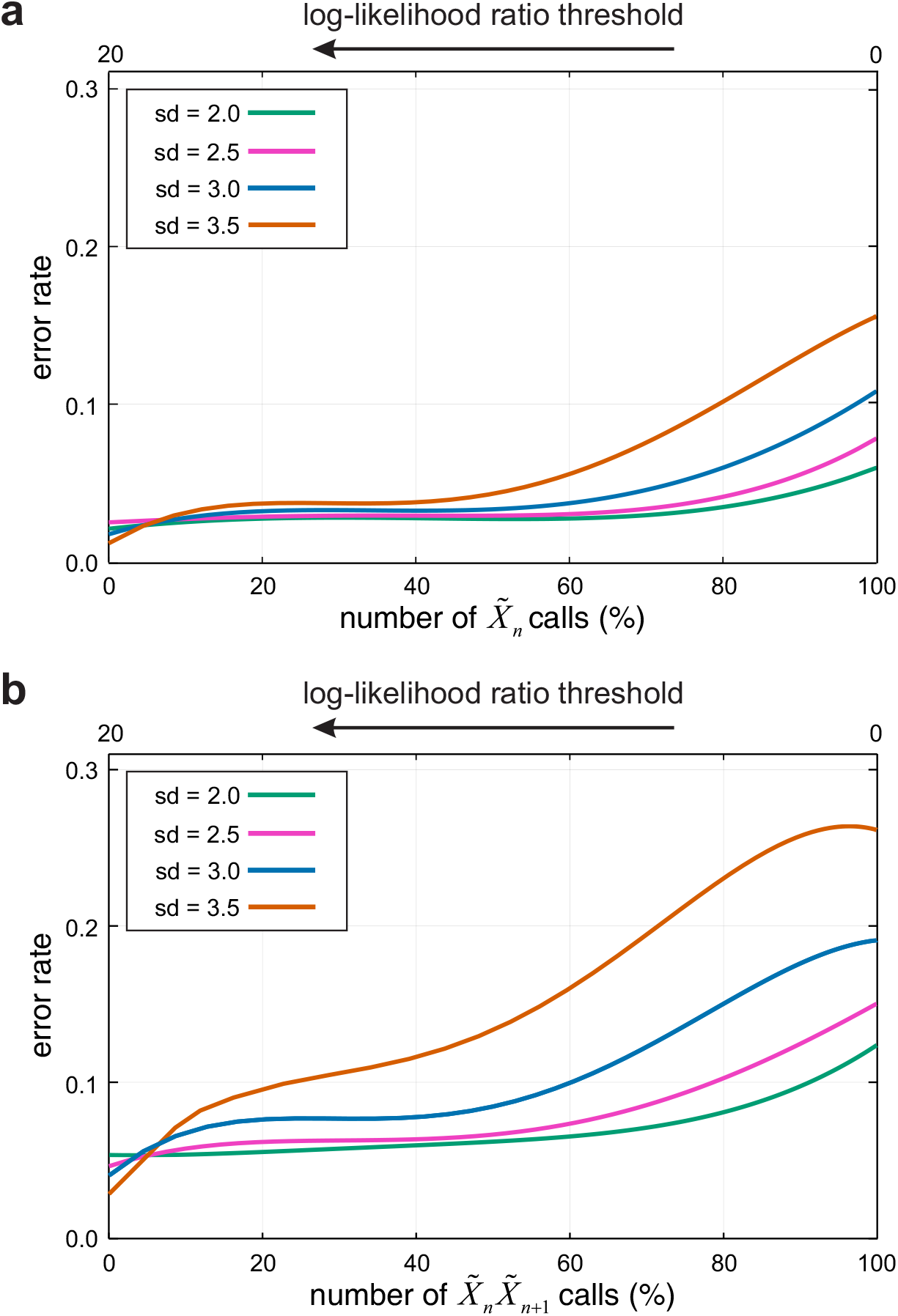
Sufficient statistics calling performance of Nanopolish. Error rates of Nanopolish in calling the true values of 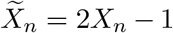, in **a**, and 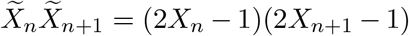, in **b**, for several signal noise levels, as a function of the log-likelihood ratio detection threshold used and the number of calls made.

On the other hand, the results depicted in Fig. 8b show that a zero log-likelihood ratio detection threshold produces an error rate in calling the true value of 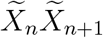 that ranges between 19% and 27% when 3 sd 3.5 and that this error remains significant even at high threshold values. This provides evidence that the sufficient statistic 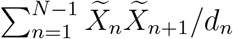, associated with the interaction parameter *γ* of the CPEL model, cannot be accurately computed from the methylation calls made by Nanopolish. If we consider also the fact that existing nanopore callers perform similarly^12^, we can conclude that the previous remark will be true for other methylation callers as well. Therefore, for reliable downstream statistical analysis of DNA methylation using nanopore data, a method must be developed, such as the one proposed here, that provides reliable estimates of methylation statistics by effectively taking into account the presence of nanopore noise.

### Performance evaluation of CPEL model estimation

We also performed simulations to evaluate the performance of the proposed EM-based maximum-likelihood method for estimating the parameters of the CPEL model from Nanopolish data, by modifying the benchmarking scheme depicted in Fig. 3 to include parameter estimation (see Fig. 9). We considered again nanopore signal noise levels with sd = 2, 2.5, 3, 3.5 and, for each standard deviation, we used the DeepSimulator to construct five sets of nanopore data with coverages *C* = 5, 10, 15, 20, 25. We did so by considering only a portion of available nanopore reads using an iterative procedure. During the first iteration, one read is picked at random from the initial pool P(0) of all available nanopore reads and two new pools P(1) and P′ (1) are formed, with the first containing the read removed from P(0) and the second containing the remaining reads. During the second iteration, one read is picked at random from P′ (1) and two new pools P(2) and P′ (2) are generated, with the first containing all reads removed from P(0) and the second containing the remaining reads. At each iteration *k*, the coverage at each CpG site is calculated as the number of reads in *P* (*k*) overlapping the CpG site, and subsequent iterations proceed until the average of all CpG coverages within Chr. 22 is less than the desired amount.

**Figure 9.**
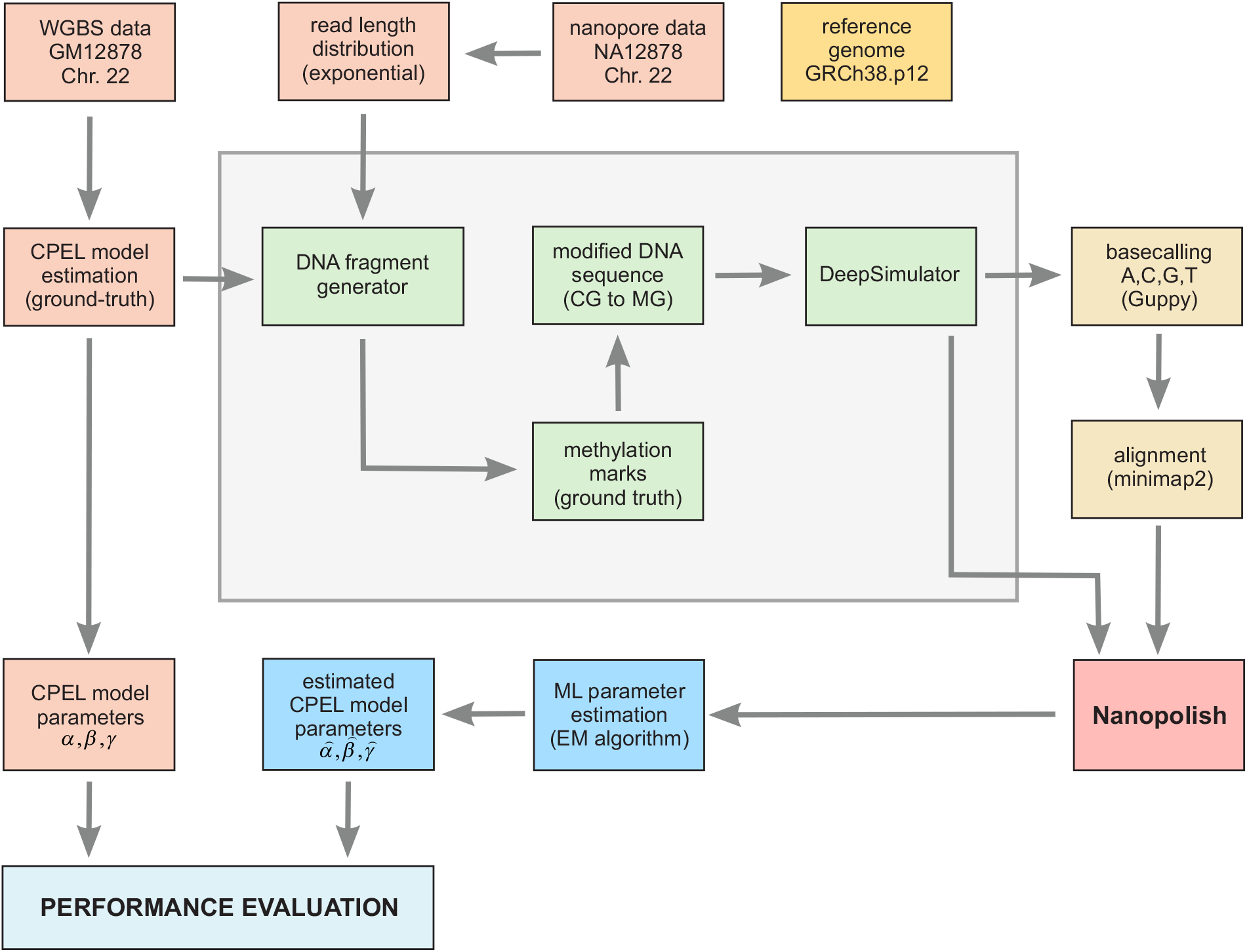
Parameter estimation evaluation scheme. Within each modeling region of Chr. 22, the values of the CPEL model parameters are determined by means of the proposed EM-based maximum-likelihood estimation approach applied on simulated Nanopolish data, which are generated by using the benchmarking scheme depicted in Fig. 3. These are then compared to ‘true’ values, which are computed by fitting the CPEL model to GM12878 WGBS data. Performance evaluation is then carried out by computing cosine similarities, boxplots, and binned probability distributions (see Figs. 10-14).

The reads in a nanopore data set with a given coverage *C* were base-called and aligned to the reference genome and subsequently used as input to Nanopolish to produce the information required for performing EM-based maximum-likelihood estimation of the *α, β*, and *γ* parameters of the CPEL model. This was performed within modeling regions that contained at least 10 CpG sites, had an average coverage of at least *C* per CG-group site, and for which methylation information was available for at least 2*/*3 of their CG-groups. To assess parameter estimation performance within a modeling region, we evaluated the closeness of estimated model parameter values 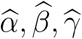 to their ‘true’ values *α, β, γ* by using cosine similarity as a measure, given by

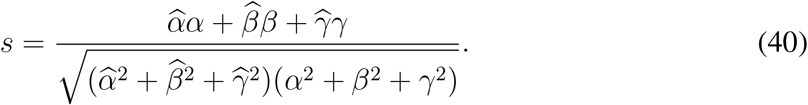

Notably, −1 ≤ *s* ≤ 1, with *s* = −1 implying maximum dissimilarity, *s* = 1 implying maximum similarity, *s* = 0 implying orthogonality or decorrelation, and in-between values indicating intermediate similarity or dissimilarity. We also evaluated, at each CpG site *n* of Chr. 22, the closeness of means E[*X*_*n*_] and pairwise correlations E[*X*_*n*_*X*_*n*+1_], predicted by the estimated CPEL models, to their true values by using the absolute error as a measure of goodness, and we did the same for the means and pairwise correlations predicted by CPEL models that were estimated directly from the methylation calls made by Nanopolish. Finally, and since empirically derived methylation statistics are commonly used in the literature, we also evaluated the correctness of estimated means and pairwise correlations that were empirically computed from the methylation calls made by Nanopolish.

The cosine similarity distributions depicted in Fig. 10 demonstrate the reliability of the proposed EM-based maximum-likelihood method for fitting the CPEL model to Nanopolish data, even at low coverage. Remarkably, the median cosine similarity values are close to 1 in all cases examined, implying that parameter estimation performs exceptionally well in at least 50% of the modeling regions considered. In addition, the estimated CPEL models predict methylation means and pairwise correlations that are mostly associated with small errors (see Figs. 11 & 12, green boxes), considering the fact that their absolute errors cannot exceed 1 since E[*X*_*n*_] = Pr[*X*_*n*_ = 1] and E[*X*_*n*_*X*_*n*+1_] = Pr[*X*_*n*_ = 1, *X*_*n*+1_ = 1], implying that 0 ≤ E[*X*_*n*_], E[*X*_*n*_*X*_*n*+1_] ≤ 1. On the other hand, and regardless of the underlying coverage, maximum-likelihood estimation of methylation means and pairwise correlations by fitting the CPEL model directly to the methylation calls made by Nanopolish consistently produced higher errors due to the presence of signal noise introduced by the nanopore sequencer. Notably, and in agreement with previous observations^13^, empirical estimation of methylation means and correlations using the methylation calls made by Nanopolish leads to substantial errors at low coverage. This is due to the fact that, in addition to not taking into account nanopore signal noise, this approach requires sufficiently large amounts of methylation data for reliable estimation, which are not available at low coverages. These results provide clear evidence that reliable downstream analysis of DNA methylation from nanopore data requires the development of a statistical approach that effectively takes into account the noise introduced on the data by the underlying nanopore chemistry.

**Figure 10.**
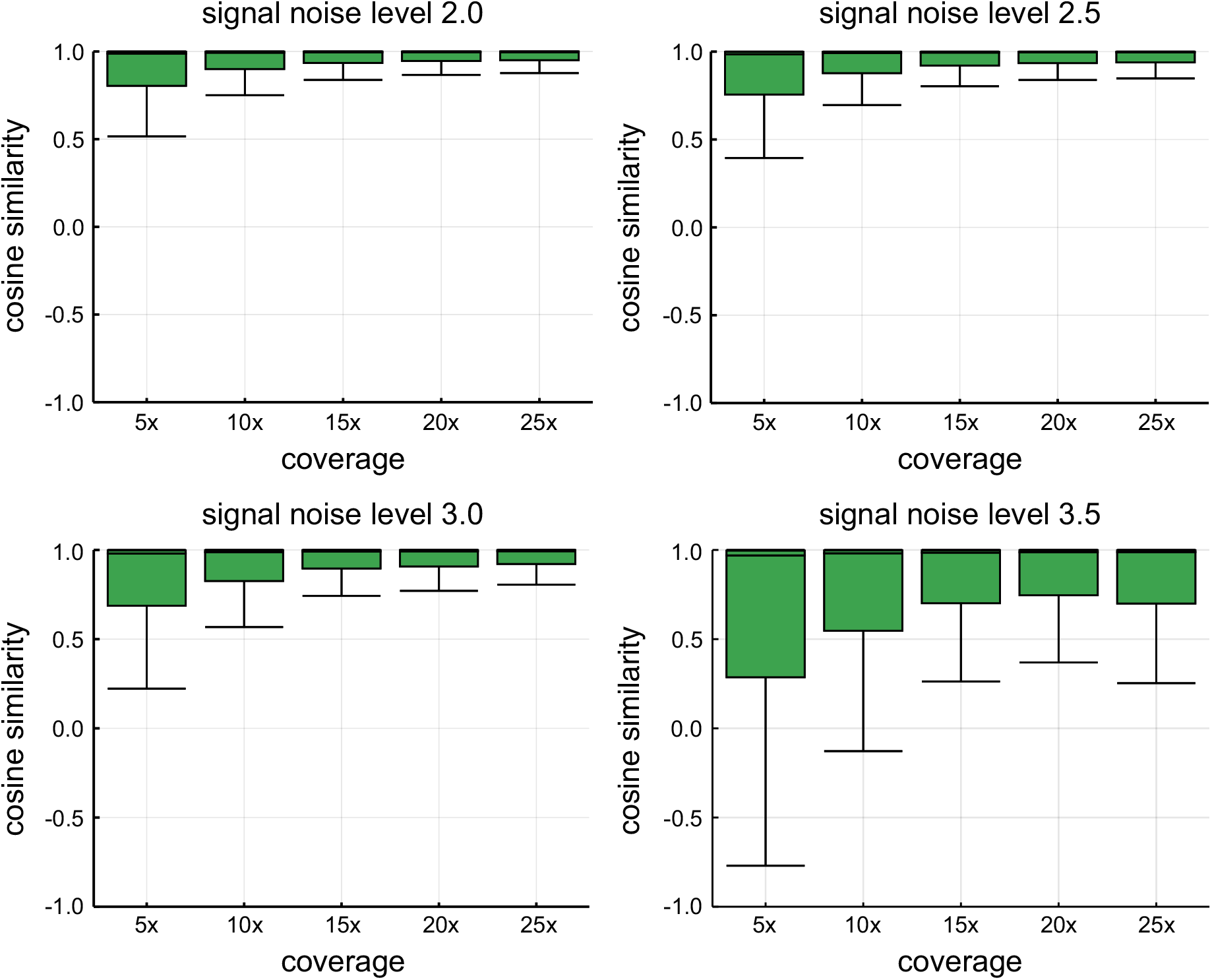
Quality of EM-based maximum-likelihood parameter estimation. Boxplots showing distributions of cosine similarities when comparing estimated to true parameter values using the simulation-based approach depicted in Fig. 9. Results are shown for signal noise levels 2, 2.5, 3, 3.5 and nanopore data with coverages 5×, 10×, 15×, 20×, and 25×. Center line of box: median value; box bounds: 25th and 75th percentiles; lower whisker: larger of minimum value and 25th percentile minus 1.5× interquartile range; upper whisker: smaller of maximum value and 75th percentile plus 1.5× interquartile range.

**Figure 11.**
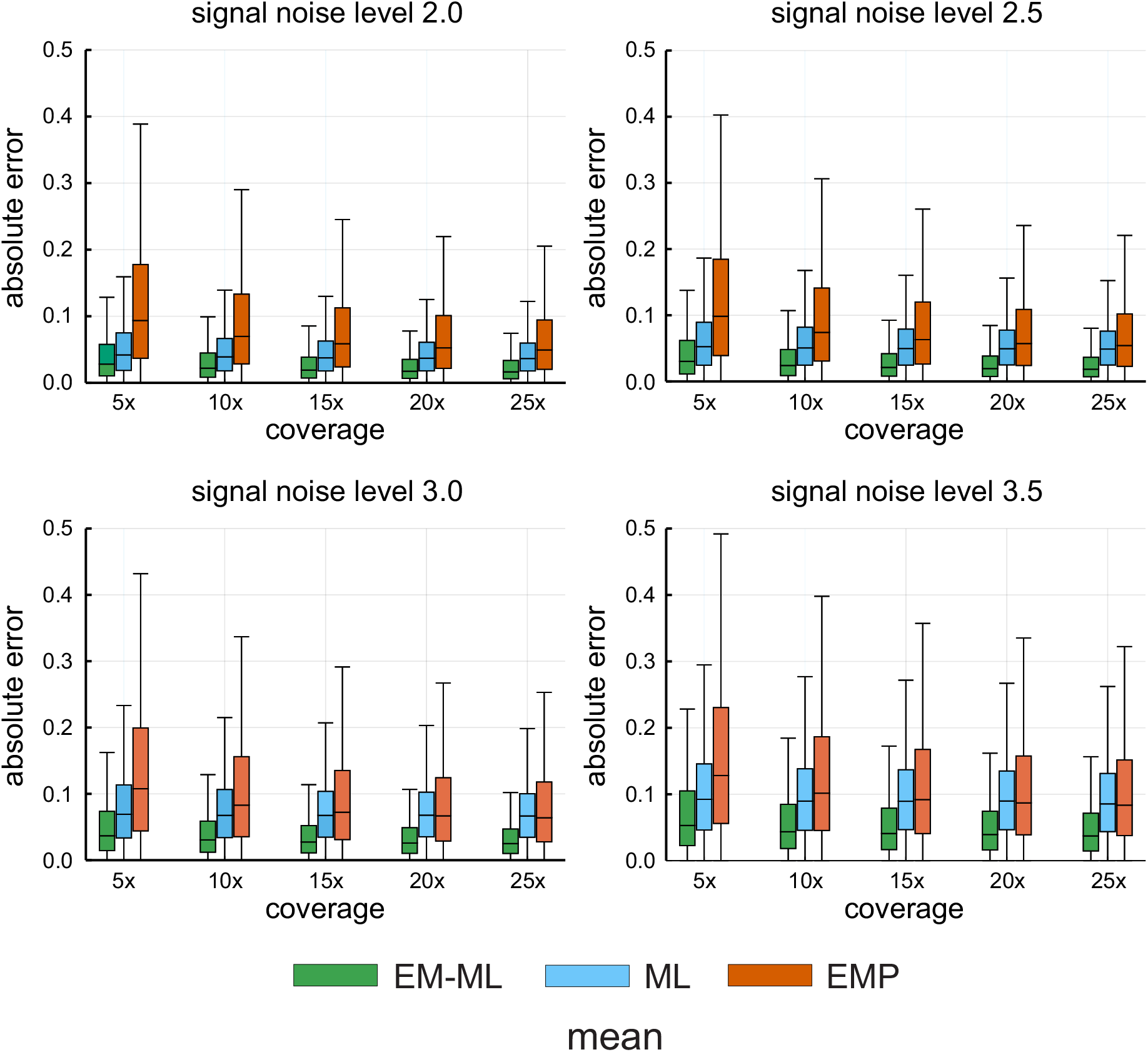
Quality of estimating methylation means. Boxplots showing distributions of absolute errors between estimated methylation means and their true values. Means were estimated by using the proposed EM-based maximum-likelihood (EM-ML) approach depicted in Fig. 9, as well as by fitting the CPEL model directly to the methylation calls made by Nanopolish using maximum-likelihood (ML) and empirically (EMP) from such calls. Results are shown for signal noise levels 2, 2.5, 3, 3.5 and nanopore data coverages 5×, 10×, 15×, 20×, 25×. Center line of box: median value; box bounds: 25th and 75th percentiles; lower whisker: larger of minimum value and 25th percentile minus 1.5× interquartile range; upper whisker: smaller of maximum value and 75th percentile plus 1.5× interquartile range.

**Figure 12.**
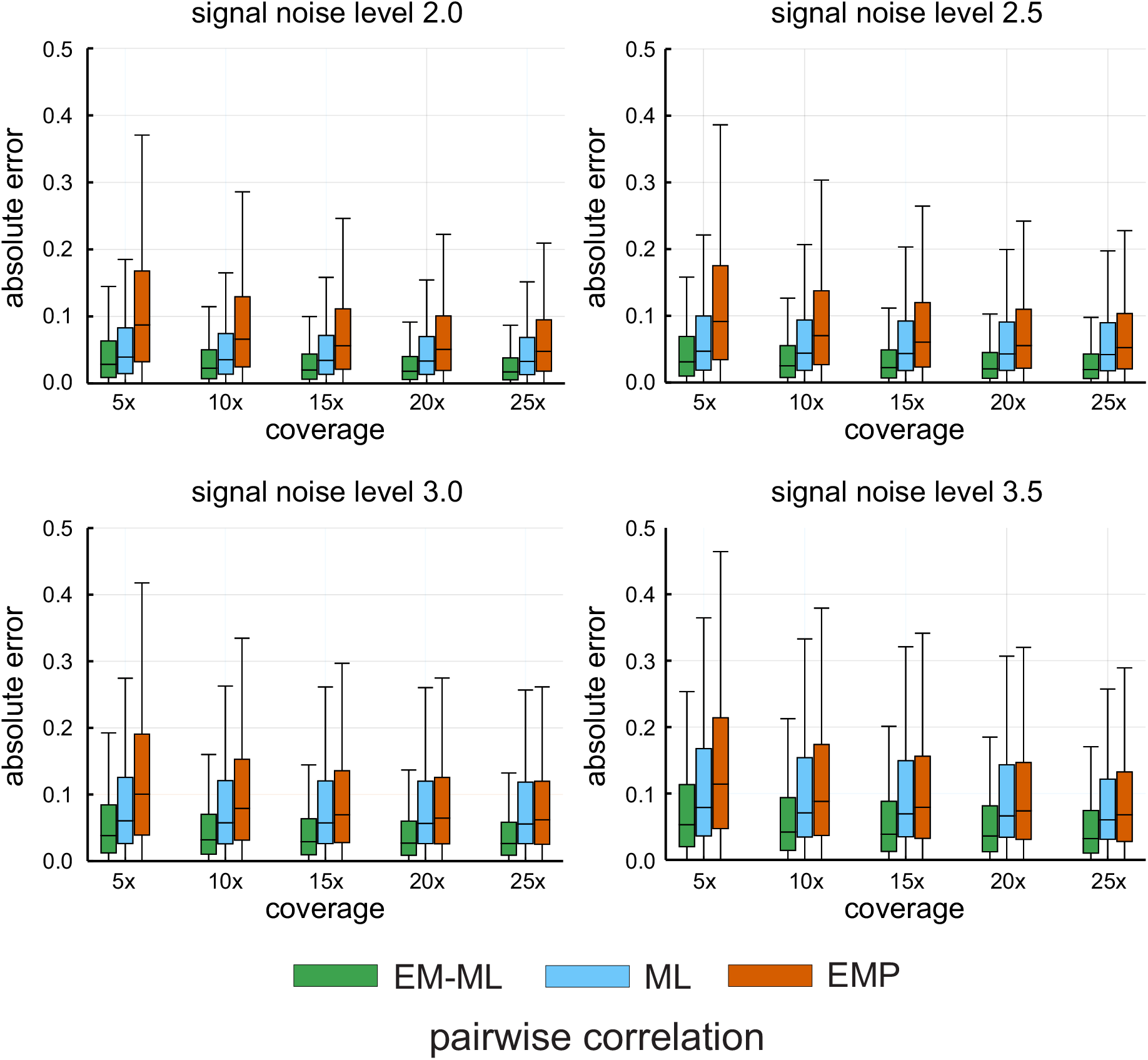
Quality of estimating pairwise correlations. Boxplots showing distributions of absolute errors between estimated pairwise correlations in methylation and their true values. Correlations were estimated by using the proposed EM-based maximum-likelihood (EM-ML) approach depicted in Fig. 9, as well as by fitting the CPEL model directly to the methylation calls made by Nanopolish using maximum-likelihood (ML) and empirically (EMP) from such calls. Results are shown for signal noise levels 2, 2.5, 3, 3.5 and nanopore data coverages 5×, 10×, 15×, 20×, 25×. Center line of box: median value; box bounds: 25th and 75th percentiles; lower whisker: larger of minimum value and 25th percentile minus 1.5× interquartile range; upper whisker: smaller of maximum value and 75th percentile plus 1.5× interquartile range.

Although Fig. 10 shows that estimation performance of the proposed EM-based maximum-likelihood approach diminishes at increasing signal noise levels, the estimated CPEL models produce reliable estimates for methylation means and pairwise correlations, especially at higher coverages (see Figs. 11 & 12). These results are also corroborated by plots of binned joint probability distributions between estimated and thur values, such as the ones depicted in Figs. 13 & 14 for signal noise level 3 and coverages 10×, 20×, which show high probabilities for most pairs of estimated vs. true values to be clustered around each plot’s diagonal. Note however that estimation of the interaction parameter *γ* of the CPEL model exhibits a skew towards higher values (see Fig. 13). We attribute this behavior to our assumption, given by Eq. (13), that the probability of finding a CG-group with variable methylation within a modeling region is negligible, which is required in order to accommodate the fact that the current version of Nanopolish assigns the same methylation state at all CpG sites within a CG-group, thus introducing an artificially higher pairwise correlation. As a consequence, modeling regions with a high proportion of CpG sites within a few CG-groups would be problematic. Nevertheless, given that almost 85% of the CG-groups in the human genome contain only one CpG site and that more than 95% of CG-groups contain at most 2 CpG sites (see Fig. 15), very few modeling regions fall into this category. As a consequence, our estimation method introduces only a slight bias in the values of the estimated pairwise correlations (see Fig. 14), which can be reduced or eliminated by a better training of Nanopolish to accommodate heterogeneous methylation within modeling regions.

**Figure 13.**
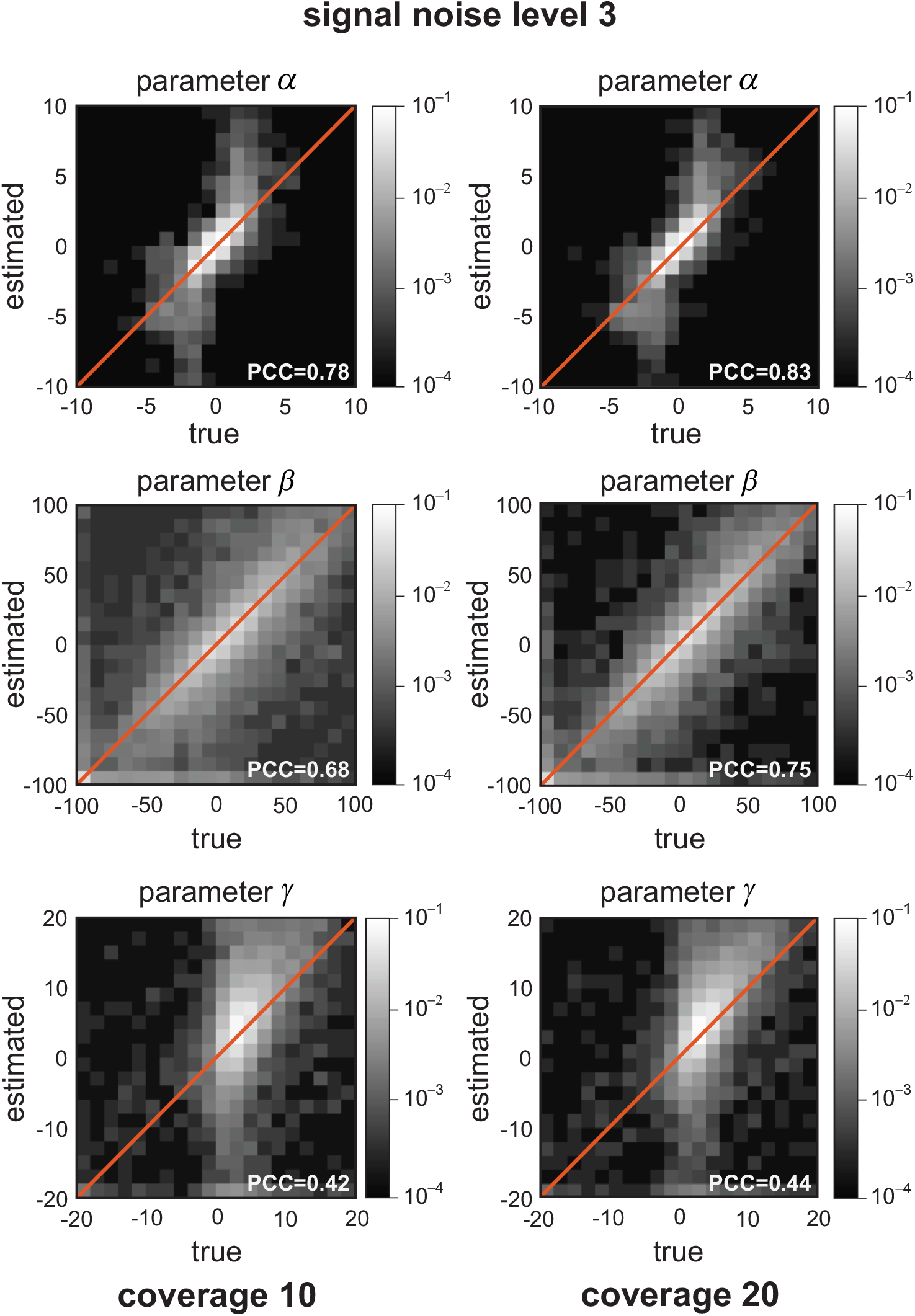
Quality of EM-based maximum-likelihood parameter estimation. Binned joined probability distributions, and associated Pearson correlation coefficient (PCC) values, between estimated CPEL parameter values and true values using the simulation-based approach depicted in Fig. 9. Results are shown for signal noise level 3 and nanopore data coverages 10× and 20×. Lighter regions indicate higher probabilities of association between estimated and true values.

**Figure 14.**
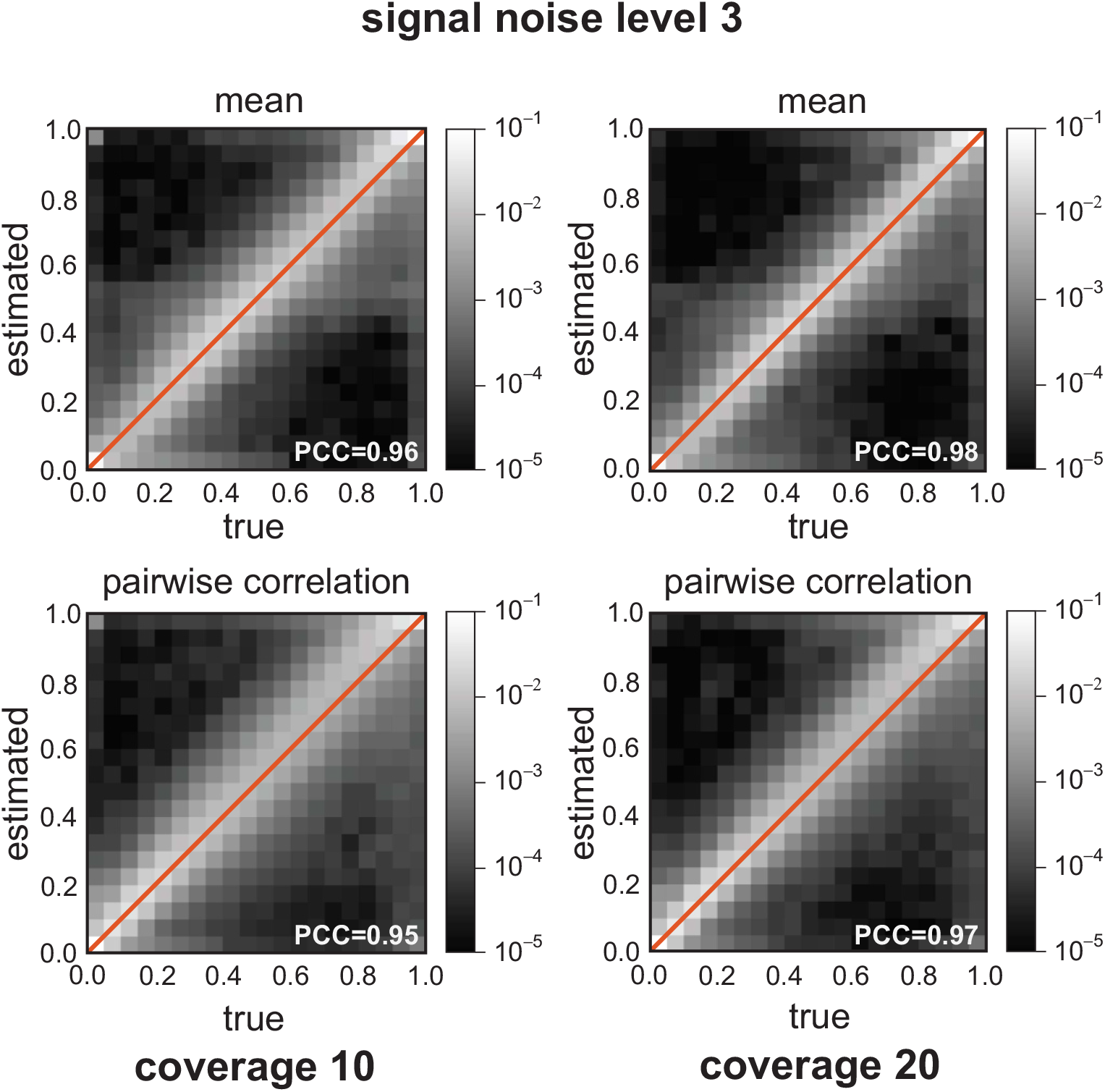
Quality of EM-based maximum-likelihood estimation of methylation statistics. Binned joined probability distributions, and associated Pearson correlation coefficient (PCC) values, between estimated means and pairwise correlations and their true values, using the simulation-based approach depicted in Fig. 9. Results are shown for signal noise level 3 and nanopore data coverages 10× and 20×. Lighter regions indicate higher probabilities of association between estimated and true values.

**Figure 15.**
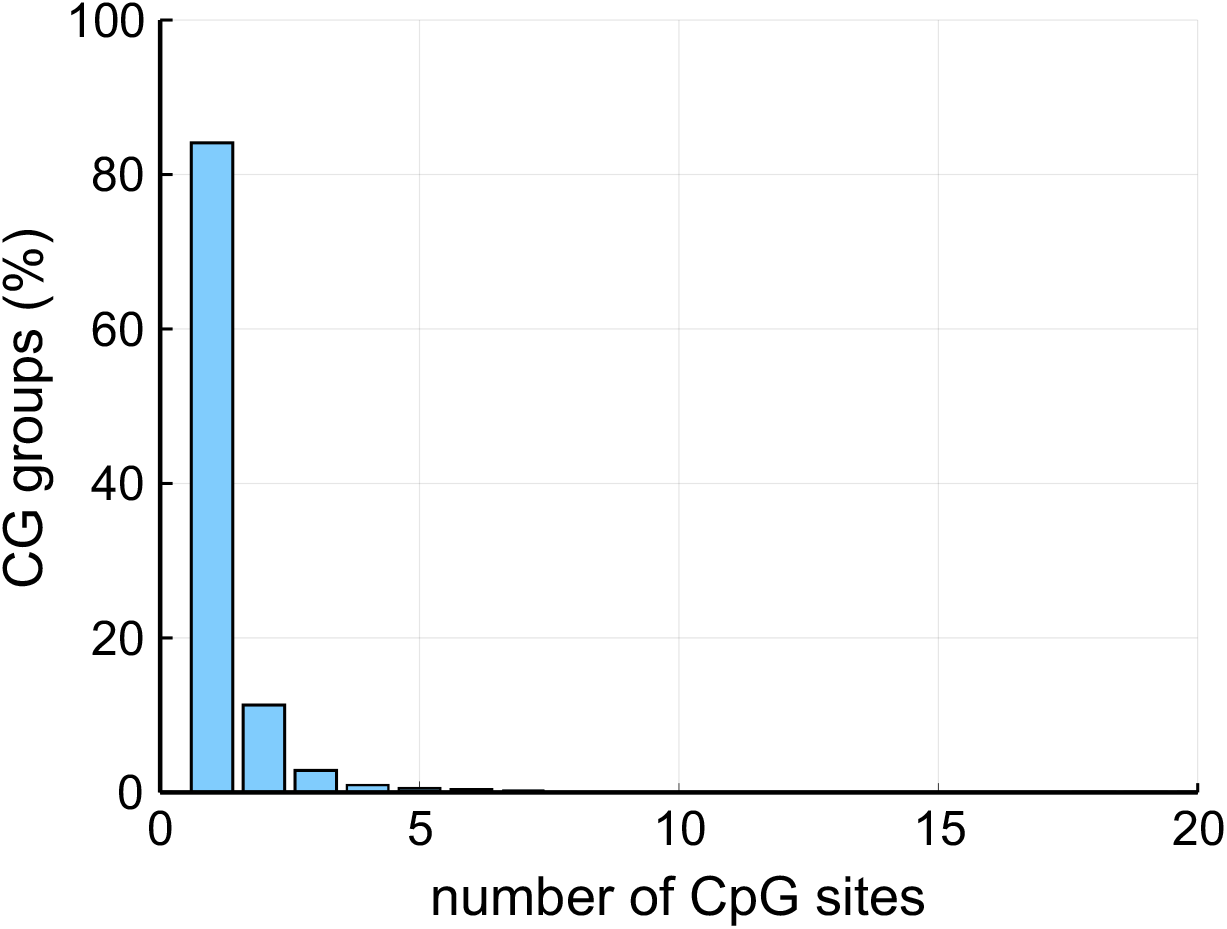
CG-group distribution. The distribution of CG-groups in the human genome in terms of CpG content.

## Supplementary Methods

### Ising computations

We here provide formulas associated with the general Ising model *p*(***x***) given by Eqs. (1)-(3), which we use for efficient computations. These formulas also apply to the CPEL model given by Eqs. (7)-(9), as well as to the reduced CPEL model given by Eqs. (20)-(22), since these models are special cases of the general Ising model. For example, the reduced CPEL model can be obtained from the general Ising model by replacing ***x***, *n*, and *N* in Eqs. (1)-(3) with ***z***, *l*, and *L*, respectively, and by setting 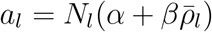 and 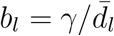.

#### Partition function

By employing the transfer matrix method^20^, we can show that

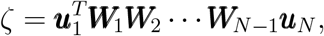

where

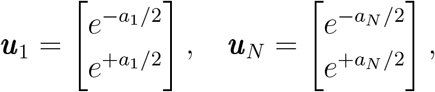

and

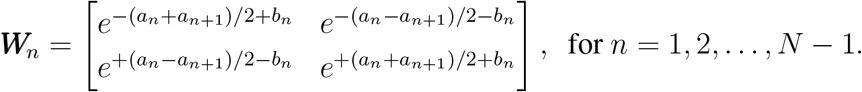

We can use this formula to compute the partition function of the general Ising model by successive vector/matrix multiplications.

#### Expectations

We compute the expectations

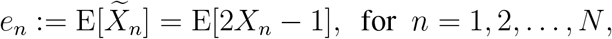

via successive vector/matrix multiplications using the following formula:

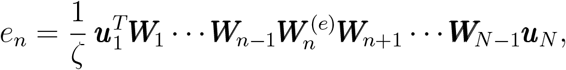

where

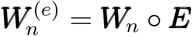

is the Hadamard product between matrices ***W***_*n*_ and

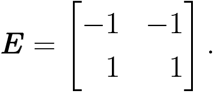

From the *e*_*n*_’s, we can then compute the expectations *µ*_*n*_ := E[*X*_*n*_] by *µ*_*n*_ = (*e*_*n*_ + 1)*/*2.

#### Correlations

The pairwise correlations

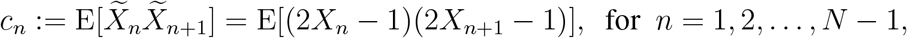

can be computed via successive vector/matrix multiplications using the following formula:

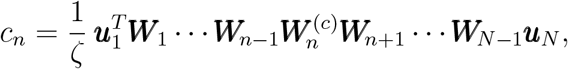

where

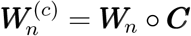

is the Hadamard product between matrices ***W***_*n*_ and

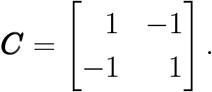

From the *c*_*n*_’s, we can then compute the pairwise correlations *r*_*n*_ := E[*X*_*n*_*X*_*n*+1_] by *r*_*n*_ = [*c*_*n*_ + 2(*µ*_*n*_ + *µ*_*n*+1_) − 1]*/*4.

#### Sampling

The general Ising model is equivalent to a first-order Markov chain with *inhomogeneous* transition probabilities. Indeed, it can be shown that

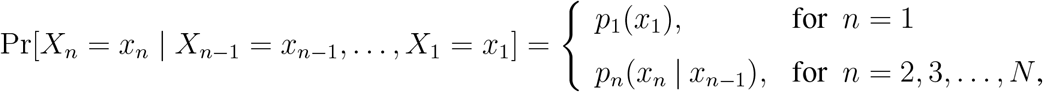

where

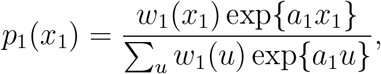

and

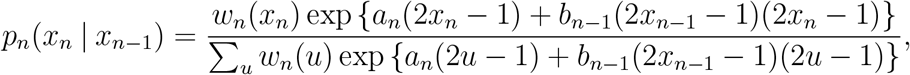

for *n* = 2, 3, …, *N*, with

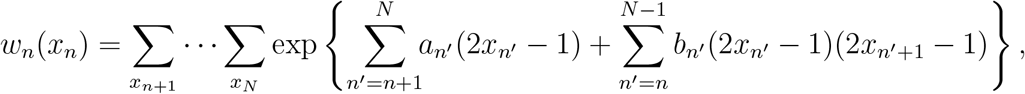

for *n* = 1, 2, …, *N* − 1, and

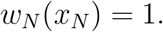

Notably, function *w*_*n*_(*x*_*n*_) can be efficiently computed using the following matrix/vector formulas:

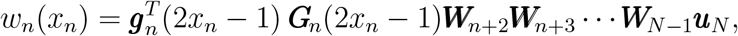

where

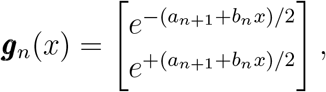

and

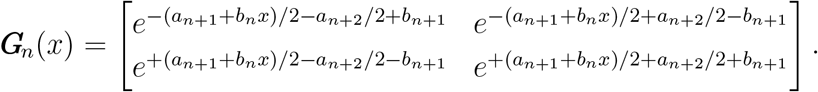

Consequently,

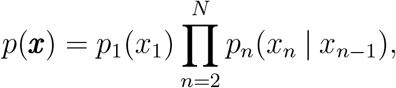

which allows us to recursively draw a sample ***x*** of the methylation state ***X*** from the Ising model *p*(***x***), by first drawing a sample *x*_1_ from the initial probability distribution *p*_1_(*x*_1_) and by sequentially drawing samples *x*_*n*_, *n* = 2, 3, …, *N*, from the transition probabilities *p*_*n*_(*x*_*n*_|*x*_*n−*1_).

### Posterior distribution of methylation indices

To compute the posterior distribution *π*(***z*** | ***y***; ***θ***) of the methylation index vector ***Z***, note that

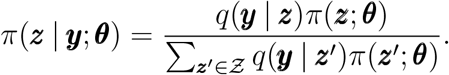

This equation, together with Eqs. (20), (21), and (27), implies that

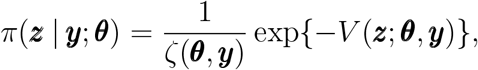

where

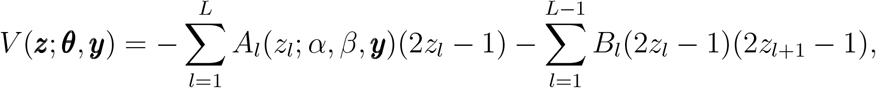

and

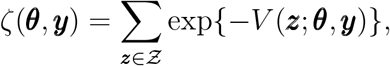

with

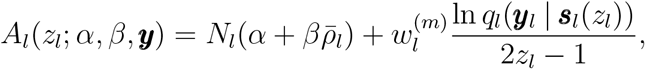

and

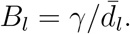

Consequently, *π*(***z***|***y***; ***θ***) is the probability distribution of an Ising model obtained by replacing ***x***, *n*, and *N* in Eqs. (1)-(3) with ***z***, *l*, and *L*, respectively, and by setting *a*_*l*_ = *A*_*l*_(*z*_*l*_; *α, β*, ***y***) and *b*_*l*_ = *B*_*l*_. In this case, computations can be preformed using the formulas provided above by replacing *N* with *L*, −*a*_*n*_ with *A*_*l*_(0; *α, β*, ***y***), +*a*_*n*_ with *A*_*l*_(1; *α, β*, ***y***), and *b*_*n*_ with *B*_*l*_.

### Data and software used

In this study, we employed the following publicly available data:

- Human reference genome built GRCh38.p12 (https://www.ncbi.nlm.nih.gov/assembly/1493941).
- GSM2308632, WGBS data (Illumina HiSeq 2500, coverage ∼100×) of human GM12878 Utah/Ceph lymphoblastoid cell line^31^ (https://www.ncbi.nlm.nih.gov/sra/SRX2157047).
- NA12878, Nanopore data (MinION R9.4 1D chemistry, coverage ∼30×) of human GM12878 Utah/Ceph lymphoblastoid cell line^8^ (https://github.com/nanopore-wgs-consortium/NA12878).

In addition, we used the following software:

- NLsolve, v4.5.0 (https://github.com/JuliaNLSolvers/NLsolve.jl).
- DeepSimulator, v1.5 (https://github.com/liyu95/DeepSimulator).
- Guppy (https://nanoporetech.com/nanopore-sequencing-data-analysis).
- Nanopolish, v0.13.2 (https://github.com/jts/nanopolish).
- minimap2, v2.17 (https://github.com/lh3/minimap2).

## Acknowledgments

We thank J. T. Simpson and W. Timp for helpful discussions.

## Funding

This work was supported by NSF Grant EFRI CEE 132452. The funders had no role in study design, data collection and analysis, decision to publish, or preparation of the manuscript.

## Notes

### Competing Interest Statement

The authors have declared no competing interest.

### Summary of Updates

Added formulas in supplementary methods for expectations and correlations in {0,1} support.

